# Dimer asymmetry in signaling of blue-light sensor histidine kinases

**DOI:** 10.1101/2025.10.24.684451

**Authors:** Vladimir Arinkin, Andreas M. Stadler, Stefanie S.M. Meier, Karl-Erich Jaeger, Andreas Möglich, Ulrich Krauss, Renu Batra-Safferling

## Abstract

Photoreceptor sensory histidine kinases (SHKs) couple light absorption to conformational changes regulating two-component signaling. Despite their importance and widespread use in optogenetics, the underlying structural signaling mechanisms remain poorly understood. We here engineered dimeric SHKs based on *Pseudomonas putida* short light-oxygen-voltage (LOV) proteins and determined their crystal structures. Regardless of illumination, the structures adopted a light-state like LOV-LOV dimer with symmetric/straight kinase modules. In contrast, small-angle X-ray scattering together with functional assays revealed pronounced light-dependent rearrangements in solution and allowed the assignment of the kinase-ON dark state to an asymmetric/kinked conformation, whereas the light state adopts a symmetric/straight structure. Comparative analyses of natural and engineered SHKs identified conserved motifs linking LOV domain rotation to kinase activity. The findings highlight the central role of dimer asymmetry and flexibility in SHK signaling, thereby not least informing the engineering of new light-responsive signaling systems.

## Introduction

Protein phosphorylation is a ubiquitous regulatory process in living organisms, central to numerous cell-signaling pathways. Mediated by kinases, it involves transferring a phosphoryl group from ATP to amino acids like serine/threonine (serine/threonine kinases), tyrosine (tyrosine kinases), or histidine (histidine kinases). In prokaryotes, sensory histidine kinases (SHKs), together with response regulators (RR), constitute so-called two-component systems (TCS). Frequently, the RR works as a transcription factor that controls gene expression in a stimulus-dependent manner. While TCS are widespread in bacteria and archaea, they have also been identified in a few eukaryotes such as plants and yeasts [1, 2]. Bacterial TCS are known to regulate a myriad of different cellular processes, including nutrient acquisition, adaptation to environmental changes such as temperature, light, osmotic pressure, and pH, as well as stress responses, cell motility, development, pathogenicity, drug resistance/ tolerance, and intercellular communication [3–11]. One generally distinguishes canonical membrane-bound SHKs [12–17] and soluble, cytoplasmic SHKs [8–11, 18–21]. SHKs are typically homodimeric and feature an N-terminal sensor module tailored to specific stimuli and a C-terminal histidine kinase (HK) effector module, which comprises dimerization/histidine phosphotransfer (DHp) and catalytic/adenosine triphosphate (ATP)-binding (CA) domains. The stimulus is detected by the sensor domain (or module) initiating autophosphorylation of the HK module, which is associated with the transfer of a phosphoryl group from ATP, bound in the CA domain, to a conserved histidine in the DHp domain. The phosphoryl group is then transferred to an aspartate residue of a cognate RR, which for many TCSs controls gene expression in phosphorylation-dependent fashion. SHKs also regulate “active” RR levels via dephosphorylation (phosphatase-activity) of the RR, forming a loop that fine-tunes cellular responses. Thus, depending on signal, one of the two activities outweighs the other, and the SHK thus serves as either a net kinase or net phosphatase on its cognate RR [22].

A subset of SHKs are photoreceptors [23] that respond to different wavelengths of the visible to near-infrared spectrum, with blue-light-sensing light, oxygen, voltage (LOV) photoreceptors and red- to near-infrared-light sensing bacteriophytochromes [8–11, 24–26] being the most widely studied systems. LOV-HKs represent interesting model systems for the study of SHK signaling, due to several reasons. First, the biological function of several LOV-HKs has been elucidated, with the systems playing an important role in bacterial physiology, ranging from regulating bacterial virulence and pathogenicity to cell-cell/cell-surface attachment and control of sulfur metabolism [8–10, 27, 28]. Secondly, HK function can be triggered by short pulses of light with high spatiotemporal resolution, enabling, in principle, the straightforward study of HK structure and function [29–31]. Finally, photoreceptor HKs including LOV-HKs and phytochrome-based systems have been widely used in optogenetics to regulate biological processes with light, such as gene expression in bacteria and yeast [32–36]. In these studies, either naturally existing photoreceptor TCS were used [36] or, the photoreceptor sensory module (e.g., the LOV sensory domain or the phytochrome photosensory core module (PCM) consisting of PAS-GAF-PHY domains), is fused to a well-characterized HK effector module, consisting of DHp and CA domains [33, 34, 37]. The first blue-light dependent engineered SHK was the chimeric protein YF1 protein [37] consisting of the N-terminal sensory domain of the *Bacillus subtilis* YtvA photoreceptor [38] and the DHp and CA domains of the FixL HK of *Bradyrhizobium japonicum* [39]. YF1 phosphorylates its cognate RR FixJ in the dark whereas upon blue-light illumination, phosphatase activity of YF1 is enhanced, thereby reducing net-kinase activity by over 1,000-fold [37]. When the YF1/FixJ TCS is used to control gene-expression in *E. coli* in a light dependent manner, this translates into high gene expression in the dark and low expression in blue light [35, 37]. YF1 function was shown to depend on the LOV photocycle [37] that involves the transient formation of a FMN-cysteinyl thioadduct [40, 41]. Formation of the light- or adduct state is commonly associated with structural changes in the LOV domain that are transmitted to fused effector domains such as e.g. the HK DHp/CA domains of YF1. In the dark, the LOV photoreceptor thermally returns to its dark state (dark recovery), which, depending on the protein, can take between seconds and hours [42, 43].

Despite fulfilling important functions in bacteria and their widespread use as optogenetic tools, high-resolution structures of full-length SHKs remain scarce, limiting our understanding of the light-dependent structural response underlying HK function and phosphorelay. In this context, soluble photoreceptor SHKs provide tractable paradigms for dissecting the structural principles of SHK signaling. At present, three structures of LOV SHKs have been reported [29–31], yet no full-length structure of a dimeric LOV-HK has been determined in both its dark and light-states, and the correlation of these states with kinase activity (i.e. ON- or OFF-states) remains elusive. Reasons for the lack of crystallization success likely include: flexibility of the HK (DHp-CA) module in either of the two states [30] and/or too fast dark recovery of the LOV-HK, which in many cases precludes crystallization of the protein in the light state [29, 43–45]. To this end, we have previously shown, that the exceedingly slow dark recovery (τ_*rec*_ ∿ 2400 min at 20°C)[46] of the short LOV protein PpSB1-LOV of *Pseudomonas putida* allows crystallization in both the dark and light state [43, 47], while its fast-recovering twin protein PpSB2-LOV (τ_*rec*_ ∿ 2 min at 20°C)[46] could only be crystallized in the dark state [48]. Given the structural similarities between the LOV domain dimer in YF1 and the PpSB1-LOV/PpSB2-LOV dimer (Figure 1), we reasoned that the replacement of the YtvA LOV domain in YF1 by the corresponding LOV domain of PpSB1-/PpSB2-LOV should enable the construction of functional light-dependent HKs, with very slow and fast dark recovery kinetics, respectively. This can yield interesting optogenetic tools for application, while at the same time increasing the chances for crystallization of both the dark and/or light state.

**Figure. 1:**
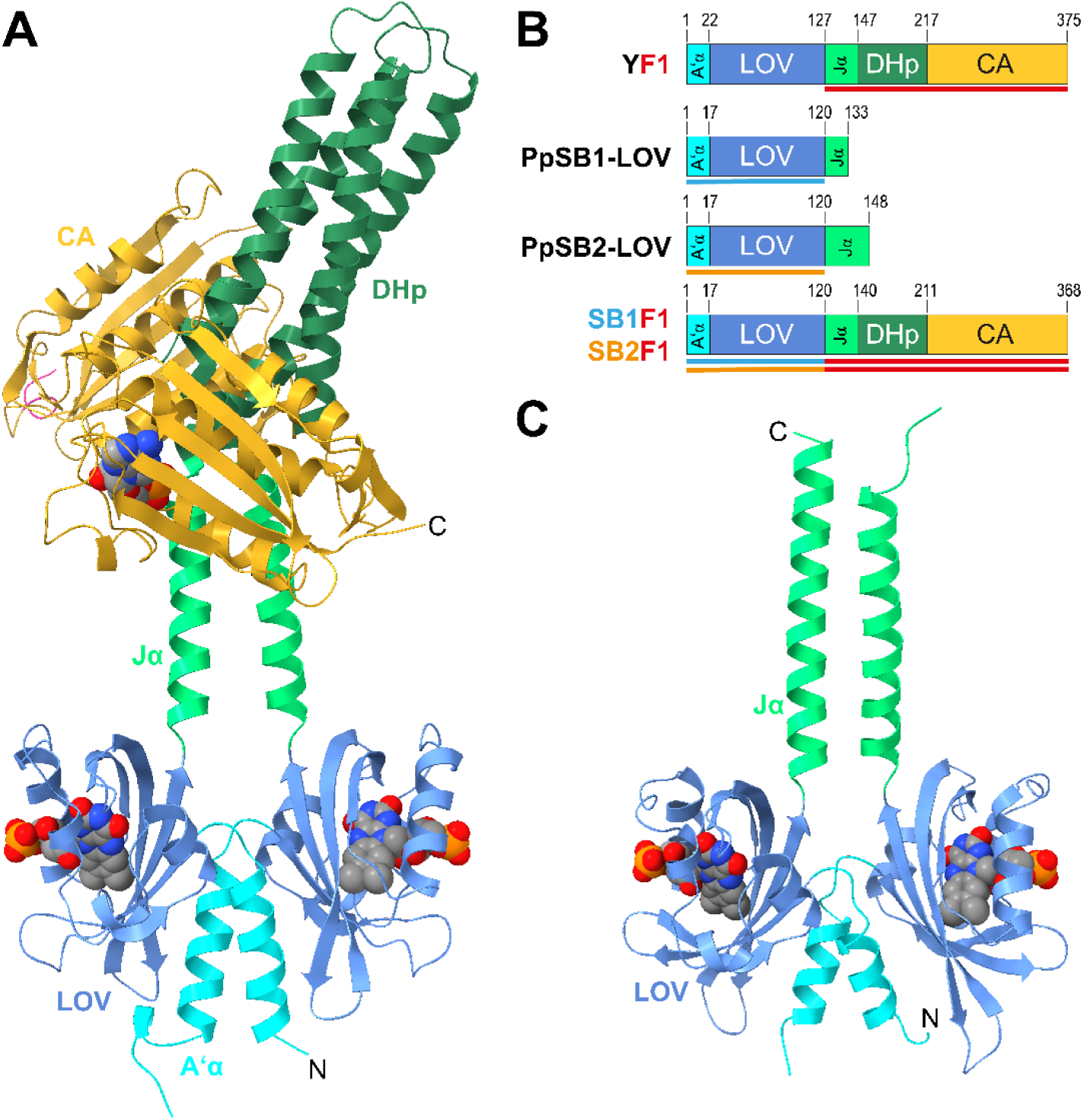
Domain architectures of the designed SHKs and structural similarity between YF1 and PpSB2-LOV. Dark-state structure (A) and multidomain architecture (B) of YF1 with the A’α (residues 1-22) in cyan, LOV domain (residues 23-127) in blue, Jα linker (residues 127-147) in light green, the rest of the DHp domain (residues 148-217) in dark green and CA domain (residues 218-375) in gold. N- and C-termini are highlighted in one of the two chains, respectively. The LOV domain FMN chromophore and the one ADP molecule bound in one YF1 CA domain are shown as VdW spheres with carbon in grey, nitrogen in blue, oxygen in red and phosphorous in orange. SB1F1 and SB2F1 were designed by exchanging the YtvA LOV domain of YF1 (residues 1-127) by the LOV domains of PpSB1-LOV and PpSB2-LOV (residues 1-120), respectively. The colored lines below the domains indicate the origin of respective domains. The dark-state structure of PpSB2-LOV (PDB-ID: 7A6P, [48])) is shown in (C). Compared to PpSB2-LOV, PpSB1-LOV has a slightly shorter C-terminal Jα helix. A multiple sequence alignment of YF1, SB1F1 and SB2F1 is shown in Supporting Figure S1.

In this study, we present the engineering and functional characterization of YF1-like sensor histidine kinases derived from *P. putida* PpSB1/PpSB2-LOV that display very different recovery kinetics lasting days for the SB1F1 protein and only minutes for SB2F1 and its slower reverting variant SB2F1-I66R. We successfully determined the dark-state structure of SB2F1, as well as both the light- and dark-state structures of the slower cycling SB2F1-I66R variant. Our findings, complemented by solution X-ray scattering studies, suggest that the activation of homodimeric LOV-SHKs is driven by light-induced transitions between symmetric and asymmetric structures, resulting in net-kinase (kinase ON) or net-phosphatase (kinase OFF) activity, respectively This structural switching suggests a mechanistic basis for how LOV-SHKs integrate light signals into opposing enzymatic outputs.

## Materials and Methods

### Cloning and site-directed mutagenesis of SB1F1, SB2F1 and SB2F1-I66R

The SB1F1 and SB2F1 fusion constructs were generated by overlap-extension PCR [49, 50]. In brief, the gene fragments encoding for the N-terminal end of the PpSB1-LOV and PpSB2-LOV photoreceptors (residues 1-120; comprising the A’α-helix and the LOV core domain; Figure 1) were amplified using the oligonucleotides SB1_ov_fw (5’-CGC GGC AGC CAT ATG ATC AAC GCG CAA TTG C -3’) / SB2_ov_fw (5’-GGC GGC CAT ATG ATC AAC GCA AAA CTC CTG CAA CTG ATG G -3’) and SB1_ov_rev (5’-GTC TGC TGG TGC TCG CTG ACG TCC TTC TGG -3’) / SB2_ov_rev (5’-GCC TGG GTC TGC TGG TGC TCT GTGACA TCG CGC TGG ATG C -3’) using a pET28a+ vector carrying the PpSB1-LOV / PpSB2-LOV encoding genes [46] as template. The gene fragment encoding the histidine kinase domain of FixL (amino acids 128–275 of YF1; including Jα helix portion of the DHp domain, Figure 1) was amplified using the oligonucleotides FixL_ov_fw (5’-GAG CAC CAG CAG ACC CAG GCG CGT CTC CAG -3’) and FixL_ov_rev (5’-GCG GCC GCA AGC TTG TCG ACT CAATTC TCG TC -3’) with pDusk [35] as template. The two gene fragments were fused in a subsequent overlap-extension PCR using the oligonucleotides SB1_ov_fw / SB2_ov_fw and FixL_ov_rev. The resulting PCR product was hydrolyzed with *Nde*I and *Sal*I and ligated into an identically digested pET28a+ vector (Novagen-Merck, Darmstadt, Germany). This cloning strategy enables the expression of the gene fusion in suitable *E. coli* expression strain under the control of the strong P*_T7_* promoter, and simultaneously results in the fusion of an N-terminally encoded hexa-histidine tag (His₆-tag; tag sequence: MGSSHHHHHHSSGLVPRGSH), which allows easy purification of the fusion protein via immobilized metal ion affinity chromatography (IMAC). SB2F1-I66R was generated by QuikChange® PCR using the oligonucleotides SB2F1-I66R _fw (5’-GGA TCA CGA CCA GCC GGG CCG TGC AAT TAT CCG -3’) and SB2F1-I66R_rev (5’-CGG ATA ATT GCA CGG CCC GGC TGGTCG TGA TCC -3’). All constructs were verified by sequencing (Microsynth-SeqLab GmbH, Göttingen, Germany).

### Heterologous gene expression

For characterization purposes, each of the above-mentioned constructs possessing an N-terminal His_6_-tag), were expressed in *E*. *coli* BL21(DE3), as described previously for the corresponding short LOV proteins [43, 47, 48]. Autoinduction (AI) media [51] was prepared with Terrific Broth medium (X972; Carl-Roth, Karlsruhe, Germany) and supplemented with 50 μM riboflavin (A6279; AppliChem GmbH, Darmstadt, Germany), 50 mg/L kanamycin and for induction, 0.5 g/L glucose, and 20 g/L lactose. Overexpression was carried out in 250 mL AI media cultures for 3 h at 37 °C with continuous shaking at 110 rpm. Afterward, the incubation temperature was reduced to 30 °C and the cells were incubated under constant agitation for another 24 h.

YF1 and FixJ were expressed in *E. coli* CmpX13 [52] as described before [29, 53]. LB medium (1 L) was inoculated with a 5 mL starter culture and grown at 37°C and 225 rpm to an OD600 of 0.6. For YF1 and FixJ the medium was supplemented with 50 µg/mL kanamycin, 50 µM riboflavin or 50 µg/mL ampicillin, respectively. Expression was induced by adding 1 mM IPTG and cultures were incubated over night at 16°C and 225 rpm.

#### Selenomethionine-labeling

The selenomethionine (SeMet) labeling protocol was adapted from Doublié (2007)[54], using the methionine auxotrophic *E. coli* strain B834 (DE3) (Novagen, Darmstadt, Germany) and SeMet medium [54] for growth. To enable adaptation to the SeMet medium, the cells were resuspended at an OD_600nm_ of 0.1 in 1 L SeMet medium supplemented with L-Methionine at a concentration of 50 mg/L (500 mL of culture in 2 L baffled Erlenmeyer flasks) and grown at 37 °C with shaking at 130 rpm until an OD_600nm_ of 1.0 – 1.5 was reached. The cells were subsequently centrifuged at 3000 *x*g for 15 min and resuspended at an OD_600nm_ of 0.35 in SeMet medium and incubated for 6 h at 37 °C and 130 rpm in order to deplete the remaining L-Methionine. Then L-Selenomethionine (purchased from Fischer Scientific, Waltham, USA) was added to a final concentration of 75 mg/L and incubated with a gradual temperature reduction to 30 °C until an OD_600nm_ of 0.6 was reached. Subsequently, gene expression was induced by addition of 1 mM IPTG. After incubation for another 4 h, the cells were harvested by centrifugation at 5000 *x*g. The wet cell pellets were either used directly or frozen at -20 °C until further use.

### Protein purification

SB1F1, SB2F1 and SB2F1-I66R were purified by using immobilized metal affinity chromatography (IMAC) as described previously [43, 46–48, 55]. The fractions containing purified protein obtained from IMAC were pooled, and the imidazole containing elution buffer was exchanged with the storage buffer [20 mm Tris pH 8.0, 40 mm NaCl], using an ÄKTA pure FPLC system (GE Healthcare, Buckinghamshire, UK) with a HiPrep 26/10 Desalting column as per standard protocol. The eluted protein fractions were pooled, supplemented with 4 mM Tris(2-carboxyethyl)phosphine (TCEP), and concentrated by ultrafiltration using Vivaspin centrifugal concentrator units (molecular mass cutoff: 10 kDa) (Sigma-Aldrich; St. Louis, MO, USA). Size-exclusion chromatography (SEC) was used as the last step in purification, and for molecular weight estimation. Superdex 200 10/300 GL and Superdex 200 Increase 10/300 GL columns (GE Healthcare; Buckinghamshire, UK) were used with an ÄKTA pure FPLC system. The standard running buffer was 20 mM Tris at pH 8.0, 40 mM NaCl. The flow rate was set to 0.5 mL/min.

YF1 was purified as described before [29]. In brief, the harvested cells were disrupted by sonification and the His-tagged YF1 was isolated via IMAC. Fractions were pooled according to yield and purity and samples were stored in 20 mM Tris-HCl, 20 mM NaCl, 10% (v/v) glycerol, pH 8.0.

FixJ was purified in two steps as described before [53]. After cell harvesting and disruption by sonification, the His_6_-SUMO-tagged FixJ was first isolated via IMAC. Fractions were pooled and the SUMO-tag was cleaved off in 50 mM Tris, 1 M NaCl, pH 8.0 overnight at 4°C by His-tagged Senp2 protease. Second, in a reverse IMAC FixJ was collected in the flow through, concentrated and stored in 20 mM Tris-HCl, 250 mM NaCl, 10 % (v/v) glycerol, pH 8.0.

### Determination of chromophore content and loading

Quantification of chromophore loading and content was performed by UV-Vis spectroscopy and high-performance liquid chromatography (HPLC) as described previously [46, 48].

### Dynamic Light Scattering

Dynamic light scattering (DLS) measurements were performed using a SpectroSize 300 instrument (Xtal Concepts; Hamburg, Germany) at 20 °C. Approximately 50 µL of protein solution (∼1 mg/mL) in the storage buffer described above was used for each measurement. Prior to analysis, all samples were centrifuged at 14,000 × g for 10 min. Scattering was recorded at a wavelength of 660 nm and an angle of 90° using a quartz glass cuvette. Measurement time was 10 s, with 20 acquisitions collected per sample. Diffusion coefficients were determined by analyzing the decay of the scattered intensity autocorrelation function, and hydrodynamic radii were calculated using the SpectroCrystal software provided by the manufacturer.

### Spectroscopic techniques

UV/Vis spectrophotometric measurements were carried out with a Shimadzu UV-1800 spectrometer (Shimadzu, Kyoto, Japan) as described previously [43, 46–48, 56]. The samples were illuminated for at least 30 s using a blue-light (λ_max_ = 450 nm) emitting LED with a radiant power of 50 mW (Luxeon Lumileds; Phillips, Aachen, Germany) to generate the respective light states. Dark recovery kinetics were then measured from the illuminated samples as described previously [43, 46–48, 56]. The absorbance recovery at 475 nm was recorded as a function of time. The absorbance data were plotted against time with the program Origin and fitted using a single exponential decay function. All measurements were done in triplicate.

### Functional characterization

The net histidine kinase activities of the SB1F1, SB2F1 and SB2F1-I66R proteins were assessed via a fluorescence anisotropy assay as reported before [53]. The binding site of phospho-FixJ (underlined) was included in (5-and-6)-carboxytetramethylrhodamine (TAMRA) labeled, double-stranded DNA (5’-GAGCGATATCT<u>TAAGGGGGGTGCCTTA</u>CGTAGAACCC-3’), which was prepared as described previously [57]. Samples were prepared under red safe light. Enzymatic measurements were conducted in 20 mM Tris-HCl, 80 mM NaCl, 2.5 mM MgCl_2_, pH 8.0 with 2.52 µM protein variant, 25.2 µM FixJ, 1.26 µM TAMRA-dsDNA and 111 µg/mL BSA [34]. Samples were transferred into a black 96-well microtiter plate (FluoroNunc) and equilibrated at 25°C for 5 min. The kinase reaction was started by adding 1 mM ATP and the TAMRA fluorescence anisotropy was monitored with a microtiter plate reader (CLARIOstar, BMG Labtech) at (540 ± 20) nm excitation and (590 ± 20) nm emission. After 25 min, the sample was ejected and illuminated with 450-nm light for 15 s, before continuing the measurement.

### Protein crystallization

The purified proteins were concentrated to ∼4 mg/mL. Crystallization experiments were performed using the vapor diffusion method in 96-well sitting-drop plates, with a drop size of 1.8 μL (0.9 μL purified protein plus 0.9 μL reservoir solution) against 70 µL of the reservoir solution. Both, SB2F1 and SB2F1-I66R proteins crystallized at 19 °C under dark conditions where the plates were kept wrapped in aluminum foil, with the reservoir containing 10% – 12% PEG 8000 (v/v), 0.2 M NaCl, 0.1 M Sodium Citrate (pH 5.8 – 6.1), 1mM ATP, 2mM MgCl_2_. Prior to cryocooling, 20 % PEG 200 (v/v) were added in small steps (∼5 %) to the drop containing crystals. All the steps were performed in the dark or under low red-light conditions. SB2F1-I66R crystallized at 19 °C under continuous light conditions where the plates were constantly illuminated with custom-made blue-light LED arrays (λ_max_ = 450 nm, Luxeon Lumileds, Phillips, Aachen, Germany), with the reservoir containing 16% – 20% PEG 3350 (v/v), 0.2 M LiSO_4_, 0.1 M Bicine (pH 9.0 – 9.3), 1mM ATP, 2mM MgCl_2_. As above, 20 % PEG 200 (v/v) were added in small steps (∼5 %) to the drop containing crystals prior to cryo-cooling.

### Single crystal microspectrometry

UV-Vis absorbance spectra of cryo-cooled crystals at 100 K were recorded in the wavelength range 250–700 nm using a microspectrometer at the beamline ID29S at ESRF (Grenoble, France) as described previously [58]. The spectra of protein crystals were measured both, before and after X-ray exposure during data collection.

### Data collection and structure determination

Single crystals were mounted in loops and flash frozen with gaseous nitrogen at a temperature of 100 K. All steps for data collection on crystals obtained in dark conditions, including data acquisition at the beamline, were performed under low red-light conditions. X-ray diffraction data at 100 K were recorded at the ESRF beamlines ID29 [59] and ID30B [60] at Grenoble, France. The respective wavelengths and corresponding detector types of each beamline are listed in Supporting Table S1. The strategy for data collection was determined using the BEST program to reduce potential radiation damage from the beam while maintaining data collection as complete as possible [61]. Data processing was performed using the XDS program [62] and AIMLESS (part of the CCP 4 package [63]). At first, we tried to obtain the initial phases by molecular replacement using the published YF1 structure (PDB ID: 4GCZ, [29]) as a search template. However, this procedure failed, despite high sequence similarity (Supporting Figure S1), and expected similar state (crystallization under dark conditions in presence of ATP/MgCl_2_). Therefore, in order to obtain initial phase information, we produced SeMet-labeled protein as mentioned above. Single-wavelength dispersion (SAD) data were collected for the crystals obtained using the SeMet-labeled protein (Supporting Table S1). Moreover, the crystals of the SeMet-labeled protein diffracted to a higher resolution. A comparison of native and SeMet-labeled protein structures showed that the incorporation of the SeMet in place of methionine did not induce any structural changes in the protein. Therefore, all presented crystal structures of SB2F1 and SB2F1-I66R were determined from crystals obtained from SeMet-labeled protein.

For the SB2F1-I66R crystals, the initial phases were determined by molecular replacement using the program MOLREP (CCP4 package)[63]. The search model was the crystal structure of SB2F1 in the dark state, obtained as part of this work. The models described were further improved with several cycles of refinement using the program PHENIX [64] and manual rebuilding using the COOT graphics program [65]. Data collection and refinement statistics are listed in Supporting Table S1.

Analysis of the SB2F1 datasets revealed that the majority of crystals belonged to the P3_1_21 space group, with unit cell dimensions of a = b ≈ 138 Å and c ≈ 97 Å (Supporting Table S1, PDB ID: 8A6X). A smaller subset of datasets (>5%) exhibited crystals in the closely related P3_2_21 space group, characterized by unit cell dimensions of a = b ≈ 138 Å and c ≈ 48 Å (Supporting Table S1, PDB ID: 8A3U). Notably, both crystal forms originated from the same crystallization drop, indicating that the variation in space group was not due to differences in crystallization conditions. Structural superposition of the two models demonstrated they are nearly identical, with a Cα RMSD of 1.09 Å over residues 1–365 (Supporting Table S2). Given that all other structures analyzed in this study adopt the P3_1_21 space group, the SB2F1 dark-state structure corresponding to PDB ID 8A6X is used for subsequent discussion and comparison.

### Small-angle X-ray scattering (SAXS)

SAXS data for all proteins were collected at beamline BM29 at the European Synchrotron Radiation Facility (ESRF) (Grenoble, France) [66, 67], as described previously [44]. The SB2F1 protein solution in buffer (20 mM Hepes pH 7.5, 150 mM NaCl, 2% Glycerol, 2 mM TCEP, 1.5 mM ATP, 3 mM MgCl_2_) was concentrated using Vivaspin 20 (Sartorius, Göttingen, Germany). The filtrate was collected and used as control during SAXS measurements. The reference measurement of the protein buffer was done before and after each protein sample. For the measurements of the protein dark-state, all sample manipulations were performed in the experimental hutch in the dark under red-light conditions. For the light-state experiments, the protein solutions were illuminated with a blue-light LED (wavelength 450 nm, radiant power 50 mW, Luxeon Lumileds, Phillips, Aachen, Germany) continuously in the sample storage position and the SAXS experiment was performed under standard light conditions. The temperature for data collection was set to 10 °C to slow down the dark recovery, thus enabling more efficient population of the light state. The X-ray wavelength used on BM29 was 0.992 Å, and the used protein concentrations were 0.64, 1.45, 2.95 and 4.18 mg/mL. For each sample, ten frames with an exposure time of 1 s each were recorded. The individual recorded frames were checked for the absence of radiation damage, and the corresponding frames were merged. The scattering contribution of the buffer was subtracted from the merged data sets of the protein solutions. The buffer-subtracted SAXS data were scaled by the measured protein concentrations. Data measured of the above-mentioned protein concentrations were merged for further data analysis. The final datasets used for evaluation were obtained by merging 0.64 mg/mL for the smaller q-range and 4.18 mg/mL data for the higher scattering vector range.

Data was analyzed and modelled using the programs available within the ATSAS software package [68]. The distance distribution function P(r) was determined using the program DATGNOM. Additionally, experimental SAXS data was compared to theoretical scattering curves of the crystal structures calculated by the program CRYSOL. *Ab initio* bead models were generated with the DAMMIN program. In both cases, 50 ab initio models were generated, aligned and averaged to the most probable model. Additional restraints were imposed to improve, for example, the connectivity and compactness of the model. The crystal structures were then aligned with the averaged model. All models were used for further averaging and filtering by DAMAVER as their normalized spatial discrepancy (NSD) were within the range of mean NSD ± 2σ (SB2F1-Dark: mean NSD = 1.406, σ = 0.324; SB2F2-Light: mean NSD = 1.354, σ = 0.367). The envelope function was determined using the SITUS package [69].

### Bioinformatic, structural analyses, modelling and graphical representation

PDB files were edited using open source Pymol version 2.6.0 (Schrödinger LCC, NY, USA)[70] or the UCSF ChimeraX software [71]. For comparative analyses of the YF1 and SB2F1 structure, we used the SB2F1 dark state structure. Interface analyses were performed using the LigPlot+ v2.2. software tool [72] and the ‘Protein interfaces, surfaces and assemblies’ service PISA at the European Bioinformatics Institute (http://www.ebi.ac.uk/pdbe/prot_int/pistart.html) [73]. Jα helix rotation/translation was analyzed by using LSQKAB [74]. The SB2F1 and YF1 structures were superposed with respect to the backbone atom positions within the first two turns of their Jα helices in both subunits (chains A and B). Next, the Jα helices of SB2F1 and YF1 were superposed with regards to the same atoms but of a single chain only (either A or B). From the resultant rotation and translation matrices, a screw-rotation axis was calculated using custom Python scripts [29]. Sequence-based coiled-coil predictions were carried out using the deep-learning based coiled-coil prediction tool CoCoNat [75] (Ref) and DeepCoil2 [76]. For structure-based coiled-coil identification, the Socket2 webservice [77] was used. The structural model of the dimeric canonical Jα coiled coil was generated using the CCBuilder 2.0 webservice (https://pragmaticproteindesign.bio.ed.ac.uk/builder/) [78]. The individual chains of resulting model were superimposed with the YF1 Jα helices (first two turns, backbone atoms only) and analyzed for clashes using ChimeraX [71]. Unless otherwise indicated, figures were generated with UCSF ChimeraX software [71] developed by the Resource for Biocomputing, Visualization, and Informatics at the University of California, San Francisco, using secondary structure assignments as given by the DSSP program [79]. UV-Vis spectra and functional assays were plotted and analyzed using the Origin 2020 software (OriginLab Corporation, Northampton, MA, USA).

### Accession numbers

Atomic coordinates and structure factors for the two SB2F1 dark-state & illuminated state, structures were deposited in the Protein Data Bank (http://www.rcsb.org) under PDB IDs 8A6X, 8A3U and 8A52, respectively. SB2F1-I66R dark & light states, were deposited under PDB IDs 8A7F and 8A7H, respectively.

## Results

### Construction of sensor histidine kinases based on PpSB1- / PpSB2-LOV modules

To construct SB1F1 and SB2F1 we replaced residues 1-127 of YF1 (comprising the A’α helix, the LOV core domain up to the YF1 DIT sequence motif (D125-I126-T127) with the corresponding parts of PpSB1-LOV (residues 1-120) and PpSB2-LOV (residues 1-120). (Figure 1, Supporting Figure S1).

The I66R mutation, which slows recovery of the parental PpSB2-LOV protein by about 7-fold [46] (marked with an asterisk in Supporting Figure S1) was introduced in SB2F1 to decelerate its recovery, with the reasoning that this could facilitate crystallization of the protein in both the dark and light states.

### SB1F1 and SB2F1 are functional light-dependent HK with disparate dark recoveries

SB1F1, SB2F1, and SB2F1 I66R were heterologously produced in *E. coli* BL21(DE3) and purified to homogeneity. UV/Vis spectra of the purified proteins (Figure 2, A-C) in their dark state showed typical LOV-protein like spectra with the main absorbance bands in the visible regions centered at 447 nm and approx. 360 nm [40–42].

**Figure 2:**
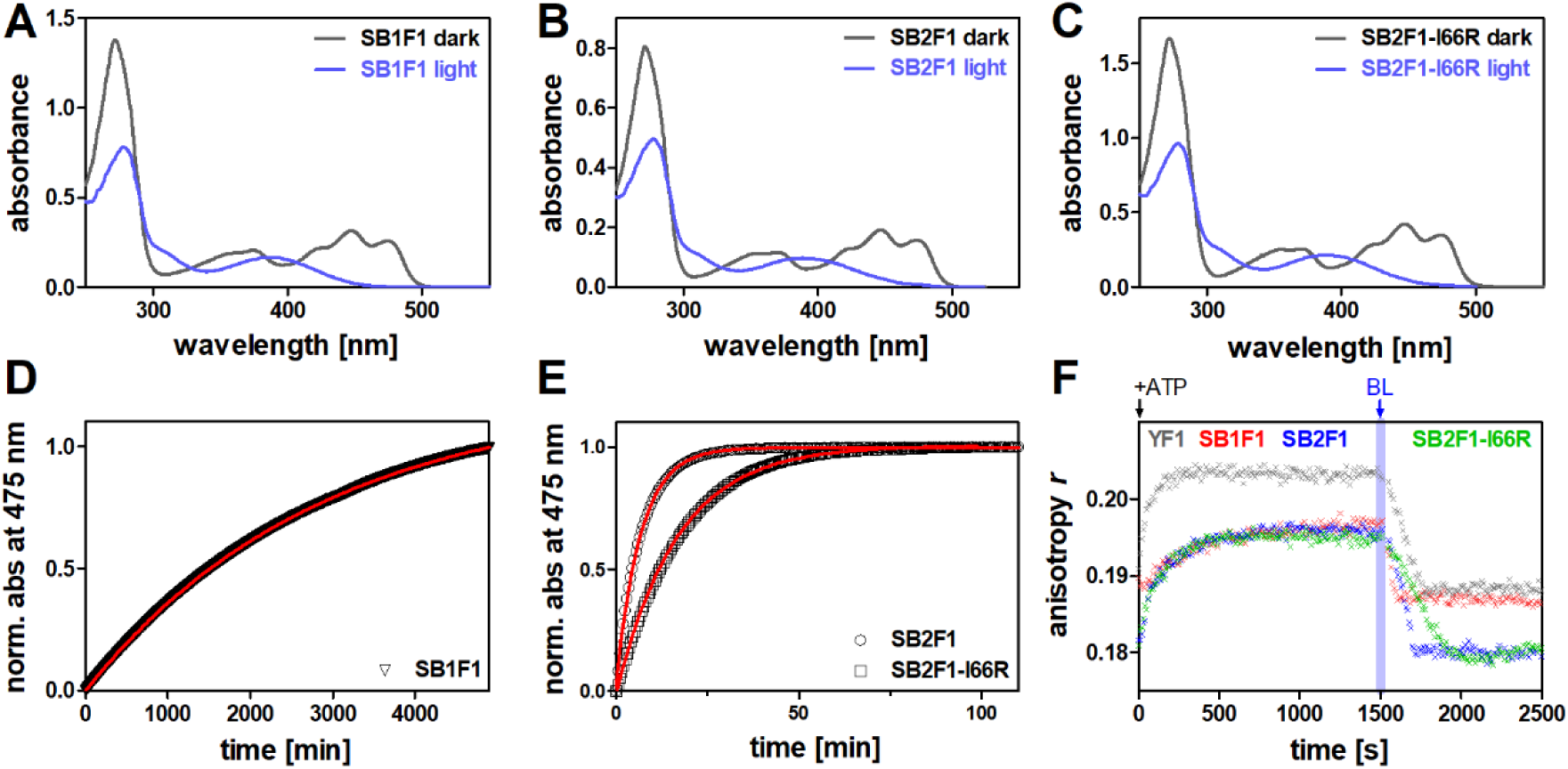
Functional characterization of SB1F1, SB2F1 and SB2F1-I66R. (A-C): Dark-(dark grey line) and light-state (light blue line) absorbance spectra of SB1F1 (A), SB2F1 (B) and SB2F1-I66R (C). Light-state spectra were recorded after illuminating the purified protein for 30 seconds with a blue LED. (D) Dark recovery time trace recorded for the SB1F1 protein after 30 seconds blue-light illumination. For SB1F1, recovery remained incomplete after 5000 min, but measurements had to be stopped due to experimental constraints. (E) Dark recovery time trace recorded for the SB2F1 and SB2F1-I66R protein after 30 seconds blue-light illumination. Data shown in D and E are normalized to aid visualization. All dark-recovery measurements were performed at 20°C. (F) Histidine-kinase activity assays. Dark equilibrated samples of SB1F1, SB2F1, SB2F1-I66R and YF1, as a control, were incubated in the dark with FixJ and a double-stranded DNA molecule containing the phospho-FixJ operator site and labeled at the 5′ end with a TAMRA fluorophore (Supporting Figure S2, A). To start the kinase reaction, ATP was added and TAMRA fluorescence anisotropy was monitored over time. An increase in anisotropy reflects kinase activity, i.e., resulting in the formation of phospho-FixJ and its binding to the DNA. Subsequent illumination with blue-light (indicated by a blue bar) caused a decrease of the anisotropy, consistent with net phosphatase activity. All experiments were performed in triplicate with consistent results (see Supporting Figure S2, B-E).

In all cases illumination with blue light resulted in the loss of the absorbance at 447 nm and the formation of a new maximum at about 390 nm, indicative of canonical LOV photochemistry involving FMN-cysteinyl adduct formation [40–42]. The recovery of the dark state of the three proteins was followed by recording the absorbance at 475 nm after illumination. As expected, from the dark-recovery kinetics of the corresponding parent LOV-proteins (PpSB1-LOV; τ_*rec*_ ∿ 2400 min, PpSB2-LOV; τ_*rec*_ ∿ 2 min, [46]), SB1F1 and SB2F1 showed very slow (SB1F1; τ_*rec*_ ∿ 2850 min) and fast (SB2F1; τ_*rec*_ = 7.0 ± 0.5 min) dark recovery kinetics, respectively. Similar to earlier studies, the I66R mutation slowed down the recovery of SB2F1 [46] by a factor of 2.5 (SB2F1-I66R; τ_*rec*_ = 17.3 ± 0.8 min).

To probe the functionality of the designed LOV-HKs, we made use of a recently reported fluorescence anisotropy assay that continuously monitors the catalytic activity and the light response of LOV-HKs like YF1 (Supporting Figure S2, A) [53]. In the dark and in the presence of ATP, YF1 phosphorylates FixJ, triggering its dimerization and binding to a 5′-tetramethylrhodamine (TAMRA)-labeled double-stranded DNA oligonucleotide that contains the FixK2 operator sequence. This binding slows DNA rotational diffusion, increasing the fluorescence anisotropy of the TAMRA label. Blue light reverses this process by increasing net phosphatase activity of YF1, leading to FixJ dephosphorylation, DNA release, and reduced anisotropy, respectively. We initially used YF1 as a control to verify the assay and for reference. As expected, upon addition of ATP, dark-adapted YF1 showed increased fluorescence anisotropy (Figure 2, F and Supporting Figure S2, B; grey symbols), whereas illumination with blue light triggered a rapid reduction of the fluorescence anisotropy, reflecting net phosphatase activity of YF1 in its light state, as previously reported [37, 53]. All LOV-HKs designed here showed very similar behavior, i.e., net kinase activity in the dark and net phosphatase activity upon blue-light illumination, albeit with slightly different kinetics and amplitudes (Figure 2, F and Supporting Figure S2, C-E, red, green and blue symbols, respectively). This, together with the spectroscopic data, reporting on LOV-photocycle integrity, demonstrates that SB1F1, SB2F1 and SB2F1-I66R are functional LOV-HKs, with very similar catalytic properties as YF1, although with altered dark-recovery kinetics.

### Crystallization and dark-state structure of SB2F1

We reasoned that the very slow dark recovery of SB1F1 (τ_*rec*_ ∿ 2850 min) could facilitate crystallization in both the light and dark states. This strategy had previously proven effective with the short PpSB1-LOV protein (τ_*rec*_ ∿ 2400 min), for which we successfully obtained crystal structures in both states [43, 47]. Unfortunately, we were unable to obtain diffraction-quality crystals for SB1F1. We therefore next focused on the fast-reverting SB2F1 protein (τ_*rec*_ = 7.0 ± 0.5 min). Native crystals were obtained using the sitting-drop vapor diffusion method (for details, see Materials and Methods). However, despite the high sequence similarity between SB2F1 and the YF1 protein (Figure 1; Supporting Figure S1), phasing attempts by molecular replacement using the YF1 model as a template were unsuccessful, already hinting at substantial structural differences. We hence obtained SeMet-substituted crystals for SAD phasing. As detailed in the Methods section, SB2F1 crystals bound to ATP/MgCl₂ were grown in the dark in space group P3₁21, and diffracted to a resolution of 2.45 Å. (Supporting Table S1). The SB2F1 dark-state structure contains two protein molecules per asymmetric unit that form a long, rod-shaped homodimer (Figure 3, A). Residues 1–365 of 368 were resolved in both monomers. In both protein chains, the LOV domain contains a non-covalently bound FMN molecule, while the CA domains are bound to ATP. Due to the relatively poor resolution in several residues of the CA domain, its refinement was restrained using the YF1 crystal structure, which has the same sequence. Superposition of the SB2F1 constituent monomers reveals moderate structural differences, reflecting a slightly asymmetry within the altogether largely C-symmetric dimer. Alignment of residues 17–138 (LOV core and DHp domain up to His138) highlights some degree of tertiary structure differences caused by DHp helix bending beyond His138 (Figure 3, B).

**Figure 3.**
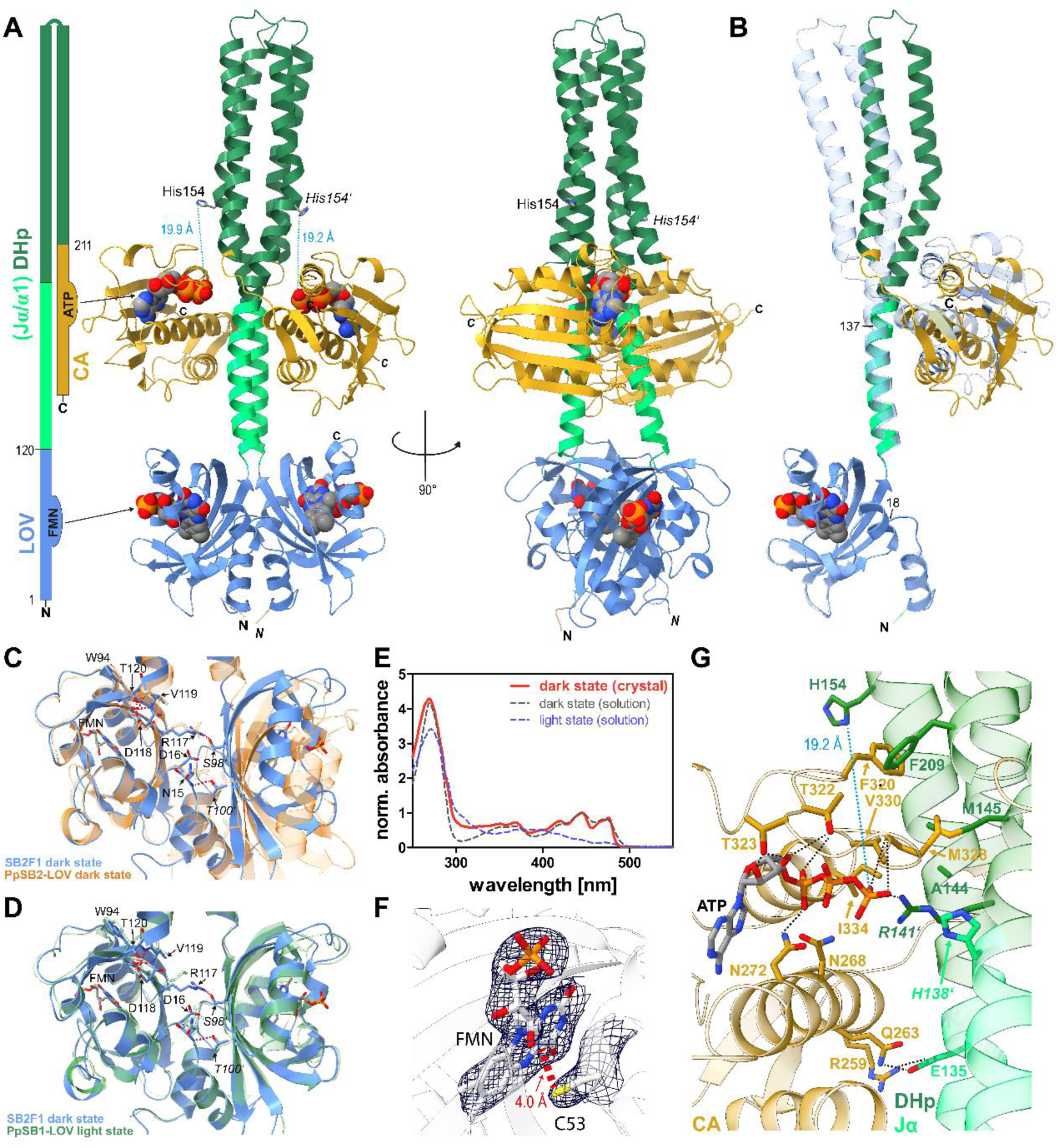
SB2F1 dark-state structure. (A) Crystal structure of SB2F1 bound to ATP, shown as a ribbon diagram with LOV (blue), DHp including Jα-helix portion (light and dark green), and CA (gold) domains. FMN and ATP are in spheres representation; His154 (phospho-acceptor) is shown as sticks. Atom colors: C–grey, O–red, N–blue, P–orange. (B) Superposition of SB2F1 chain A (colored as in (A) with chain B (transparent light blue), aligned over residues 17–138 to highlight divergence beyond the DHp domain. (C, D) Superpositions of the SB2F1 dark-state LOV dimer (blue) with the dark-state PpSB2-LOV (orange, C) and light-state PpSB1-LOV (green, D) dimers. FMN is shown as sticks; dimers were aligned over residues 1–120. Key inter- and intrachain interactions are shown in stick representation with C–light blue, O–red, N–blue, P–orange; for SB2F1 and light-orange and light green for the PpSB2-LOV dark-state (C) and PpSB1-LOV light state (D) respectively. (E) Single-crystal microspectrometry data of dark-grown SB2F1 crystals (red) compared to SB2F1 in solution (dark-state: dashed grey; light-state: dashed blue), normalized at 450 nm. (F) σ-A weighted 2mFo-DFc electron density map of the FMN and residue cysteine 53 of SB2F1 contoured at 1.0 σ. (G) SB2F1 dark-state structure as in (A), highlighting CA and DHp residues interacting with ATP and mediating interdomain contacts. H-bonds (<3.2 Å) shown as black dashed lines; His154^…^ATP-γ-P distance shown as cyan dashed line in (A) and (G). Interacting residues from the other chain are italicized throughout.

Interestingly, superposition of the dark-state dimer of the short LOV protein PpSB2 on the SB2F1 dark-state LOV-LOV dimer results in a poor alignment (Cα RMSD over residues 1-118; 3.23 Å) (Figure 3, C). In contrast, superposition with the light-state structure of PpSB1-LOV yields a significantly better fit (Cα RMSD over residues 1-118: 1.05 Å) (Figure 3, D) (Supporting Table S2). Hence in terms of the overall LOV-LOV dimer, the SB2F1 dark-state structure seems to be much closer to a light-state structure (here represented by the light-state structure of PpSB1-LOV, PDB-ID: 3SW1 [43]; since no PpSB2-LOV light-state structure is available), than to the corresponding PpSB2-LOV dark-state dimer (PDB-ID: 7A6P, [48]). This apparent conundrum is also reflected in intra- and interchain interactions centered around the LOV-LOV dimer interface, that are relevant for signaling.

In SB2F1 those interactions include interchain contacts between N15**^…^***T100’*, R117**^…^***S98’* (interacting residues from the symmetric dimer chain are italicized and marked with prime notation) and intrachain contacts between residues of the DVT motif and W94 (D118**^…^**T120, V119**^…^**W94) (compare Figure 3, A and B). A similar interaction pattern is present in the PpSB1-LOV light-state structure and is seen in molecular-dynamics simulations [80, 81]. In contrast, the pattern is changed in the corresponding dark-state structure of PpSB1-LOV that displays an N15**^…^***S98’* interaction and an R117**^…^**D16 interaction (not shown), while the interactions among the residues of the DVT motif remain undisturbed. To verify that the crystals grown under dark conditions remained in the dark state during preparation, single-crystal microspectrometry was performed on the dark-grown SB2F1 crystals immediately prior to X-ray diffraction data collection. The results clearly show a canonical dark-state spectrum for the crystal, consistent with the spectrum observed in solution (Figure 3, E). Additionally, in the corresponding SB2F1 dark-state 2mFo-DFc electron density map no continuous density is observed between the FMN-C4a atom and the C53-SG atom, with C53 adopting a single conformation pointing away from the FMN-C4a atom (FMN-C4a**^…^**SG-C53; 4.0 Å) that is commonly associated with the structures of dark-adapted LOV receptors [47, 82–84] (Figure 3, F). Thus, both single-crystal microspectrometry and the electron density around the FMN/C53 region clearly indicate that the FMN molecules in the individual LOV domains are in a dark-adapted state, with no covalent bond between the FMN-C4a atom and C53, although the LOV-LOV dimer arrangement within SB2F1 more closely resembles the conformation seen for the parental PpSB1-LOV light-state dimer (Figure 3, D).

In both CA domains of the SB2F1 dimer (Figure 3, A and G), ATP can be fitted into the electron density map (Supporting Figure S3). The distance between the phospho-accepting His154 in the DHp domain and the ATP γ-phosphate is 19.2 Å in chain A and 19.9 Å in chain B (Figure 3, A), thus too far to enable phosphotransfer (in *cis*) between ATP in the CA domain and His154. Therefore, the HK domain structure likely corresponds to a ‘kinase OFF’ state. Figure 3, G shows a closeup view of the ATP-binding CA domain and the CA-DHp interface of SB2F1, highlighting key interactions between residues within the CA and DHp domains that interact with the ATP molecule. The ATP molecule is hereby coordinated by residues within the conserved N, F, G1, and G2 sequence regions [85], involving H-bonding interactions to the side chains of N272, T322, T323 and the M328, G329 backbone. In the ATP binding pocket of the SB2F1 CA domain, there is continuous electron density between N268, N272 and the phosphate groups of ATP, even at high σ contouring of the 2mFo-DFc electron-density maps (Supporting Figure S3, A). The mFo-DFc map, contoured at 3σ, shows some additional density not covered by the model (Supporting Figure S3, A). Superposition with the structurally similar kinase domain, WalK (Supporting Figure S3, B) [86], showed that WalK with bound ATP has a magnesium ion located approximately in this continuous density. Although the Mg²⁺ ion was not modeled in the SB2F1 structure due to the low resolution of the CA domain maps, the strong alignment of ATP and the kinase domains between the two structures suggests that an Mg²⁺ ion likely contributes to ATP binding and catalysis in SB2F1 as well. The interaction between the CA and DHp domains is mostly mediated by hydrophobic interactions (F320 ^…^ F209, M328 ^…^ M145, and centered around I334, V330, A144), hydrogen bonds (E135 ^…^ Q263) and salt bridges (E135 ^…^ R259). Residue R141 on the DHp domain interacts with the ATP γ-phosphate.

### Illumination of dark-grown SB2F1 crystals only results in local photoactivation

The crystallization of SB2F1 under continuous blue-light illumination yielded crystals, which, however, diffracted only to a very low resolution (lower than 10 Å). Therefore, SB2F1 crystals grown in the dark with ATP were illuminated with blue light prior data collection to assess light-induced structural changes; this structure is referred to as SB2F1 “illuminated-state”. Nonetheless, due to crystal lattice constraints, it does not reflect the “true” light state, as previously observed for other LOV proteins [47]. Indeed, when we compare the structures of dark-grown SB2F1 and “illuminated” SB2F1, only minor global differences can be observed (Supporting Figure S4) (Cα RMSD of 0.64 Å superimposed over LOV-LOV dimer). When the two structures are superimposed via chain A of the LOV-LOV dimer, the differences become visible (Supporting Figure S4, A). These include a slight rotation of the chain B LOV dimer relative to chain A accompanied by a minor displacement of the DHp domains which is amplified to both CA domains (Supporting Figure S4, A), corroborated by increased residue-wise RMSDs for the CA domains (Supporting Figure S4, B). While single-crystal microspectrometry data clearly verifies light-state like UV/Vis spectra for the illuminated crystal (Supporting Figure S4, C), the corresponding 2mFo-DFc electron density map did not show any continuous electron density between FMN-C4a and C53-SG (Supporting Figure S4, D), which might be attributed to X-ray induced radiation damage, as described for other LOV protein structures [43, 87–89]. Local photoactivation in the crystal is also indicated by rotation of C53 to fully occupy the position atop the flavin C4a atom in one of the chains, accompanied by a flipping of Q116, which are hallmarks of LOV photoactivation [53, 90, 91] (Supporting Figure S4, D).

### Dark and light-state SHK structures can be obtained by crystallization of the slower cycling SB2F1-I66R variant

Next, to enable crystallization of SB2F1 under continuous blue-light illumination, the above described SB2F1-I66R variant was used, which possesses a prolonged adduct-state lifetime of τ_*rec*_ = 17.3 ± 0.8 min (Figure 2, E) as compared to SB2F1 (SB2F1; τ_*rec*_ = 7.0 ± 0.5 min), potentially enhancing protein crystallization in the light state, while maintaining SHK functionality (Figure 2, F). SB2F1-I66R with bound ATP, could be crystallized in the dark and under continuous blue-light illumination (light state) (Supporting Table S1). For both conditions, crystals were obtained in space group P3_1_ 2 1 diffracting up to 2.71 Å and 3.145 Å, respectively. Since we observed a light-state like LOV-LOV dimer arrangement in the SB2F1 dark-state structure (see above), we first assessed whether the SB2F1-I66R dark-state structure also adopts a light-state like LOV-LOV dimer. As observed before for SB2F1, the superposition of the SB2F1-I66R dark-state structure with the PpSB2-LOV dark-state dimer, yields a much worse fit (Cα RMSD over residues 1-118; 3.23 Å) than the superposition with the PpSB1-LOV light-state dimer (Cα RMSD over residues 1-119; 1.05 Å) (Supporting Figure S5, A and B; Supporting Table S2). Superposition of the SB2F1-I66R light-state structure with the PpSB1-LOV light-state dimer yields a low Cα RMSD of 0.86 Å (over residues 1-118), while the superposition with the PpSB2-LOV dark state dimer yields a Cα RMSD of 3.09 Å (over residues 1-118), indicating that the overall arrangement of the LOV-LOV dimer does not change dramatically in the SB2F1-I66R light-state structure as compared to the dark state. This can also be seen, when the SB2F1-I66R dark and light-state structures are superposed via chain A of the dark state dimer (Figure 4, A). A more detailed analysis of the structural differences can be found in the Supporting Information (Supporting Results 1, Supporting Figure S6). In order to verify that the structure obtained from light-grown crystals represents a light-state conformation (inferred from FMN-Cys adduct formation), we recorded single-crystal microspectrometry data for the light-grown SB2F1-I66R crystal. The results clearly confirm adduct formation in the crystal (Figure 4, B). We therefore modelled the FMN-C4a atom as *sp*3 hybridized, as also reported in our previous PpSB1-LOV and SBW25-LOV light-state structures [43, 55]. However, although the corresponding 2mFo-DFc electron shows continuous electron density between FMN-C4a and C53-SG (Figure 4, C), and the Cys53 side chain has moved to a position directly on top of the FMN-C4a atom, the FMN-C4a ^…^ Cys53-SG distance is with 2.7 Å, too long for a covalent bond. These observations may be attributed to X-ray induced radiation damage, as reported also for other LOV protein crystal structures [43, 87–89].

**Figure 4:**
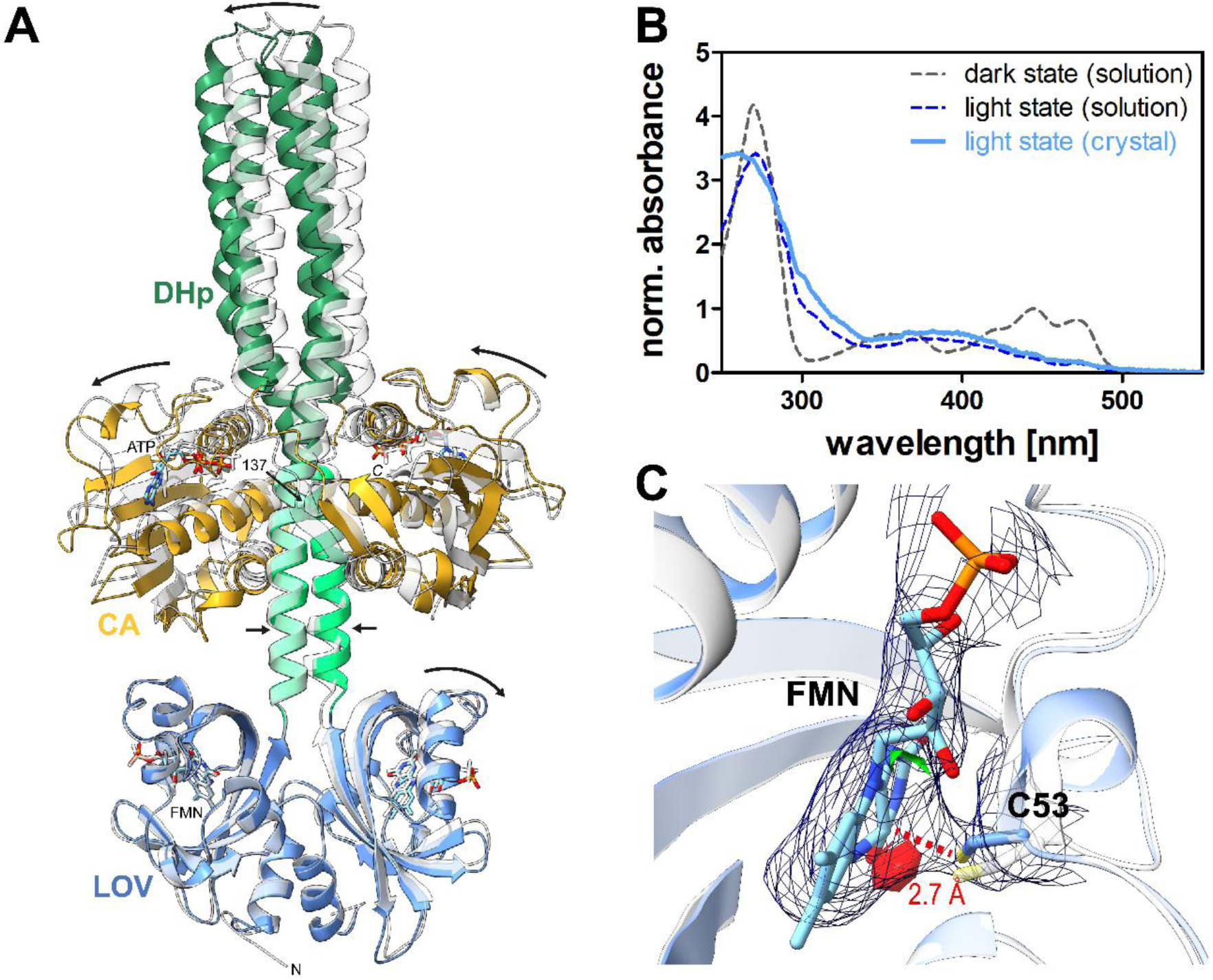
Comparison of SB2F1-I66R dark- and SB2F1-I66R light-state structure. (A) Crystal structures of the SB2F1-I66R dark-state structure (transparent grey ribbon) superimposed with the SB2F1-I66R light-state structure (solid ribbon) with the domains colored as described in Figure 3. FMN and ATP in stick representation with C–grey, O–red, N–blue and P–orange in the light-state structure and are kept grey transparent in for the dark state structures. Structures superimposed via chain A of the LOV-LOV dimer. Arrows mark the direction of dark-to-light state movements. (B) Single-crystal microspectrometry data of SB2F1-I66R light-state crystal (solid light blue line) in comparison to SB2F1 solution dark-(dashed grey line) and light-state spectra (dashed blue line). All spectra were normalized to the 450 nm (dark state) and 390 nm (light state) absorbance band, respectively. (C) 2mFo-DFc electron density map of the FMN chromophore and Cys53 of the SB2F1-I66R light-state structure contoured at 1.0 σ. The corresponding mFo-DFc map is shown as green (3.0 sigma) and red (-3.0 sigma) colored surface 2 Å from the FMN and C53.

### Small-angle X-ray scattering studies in solution reveal light-dependent structural changes of the designed SHKs

Given the above ambiguities in assigning a functional state to both the LOV-LOV dimer and the HK module in our structures, we next recorded solution small-angle X-ray scattering (SAXS) data for SB2F1 in the presence of ATP, both in darkness and after illumination with blue light (Supplementary Table S3).

The corresponding SAXS scattering data (Figure 5, A, B) indicates that the samples are homogeneous and non-aggregating. Independently, SAXS, analytical size exclusion chromatography (SEC) and DLS measurements confirmed that SB2F1 (with bound ATP) is a dimer in both the dark and light state (Supplementary Table S3), with SEC overestimating the molecular weight, probably due to the elongated, non-spherical shape, of the molecule. Guinier analysis (Figure 5, B) suggests that the dark state (*R_g_* = 39.6 ± 0.1 Å) is slightly more extended compared to the light state (*R_g_* = 37.3 ± 0.1 Å) (Supplementary Table S3). Large-scale conformational differences are clearly visible from the Kratky plot and the Pair distribution *P(r)* function, with a Kratky plot indicative of the multidomain nature of the protein. In order to assess how the here determined SB2F1 crystal-structures differ from those in solution, the theoretical scattering profiles of all here determined SB2F1 crystal structures were compared to experimental data.

**Figure 5:**
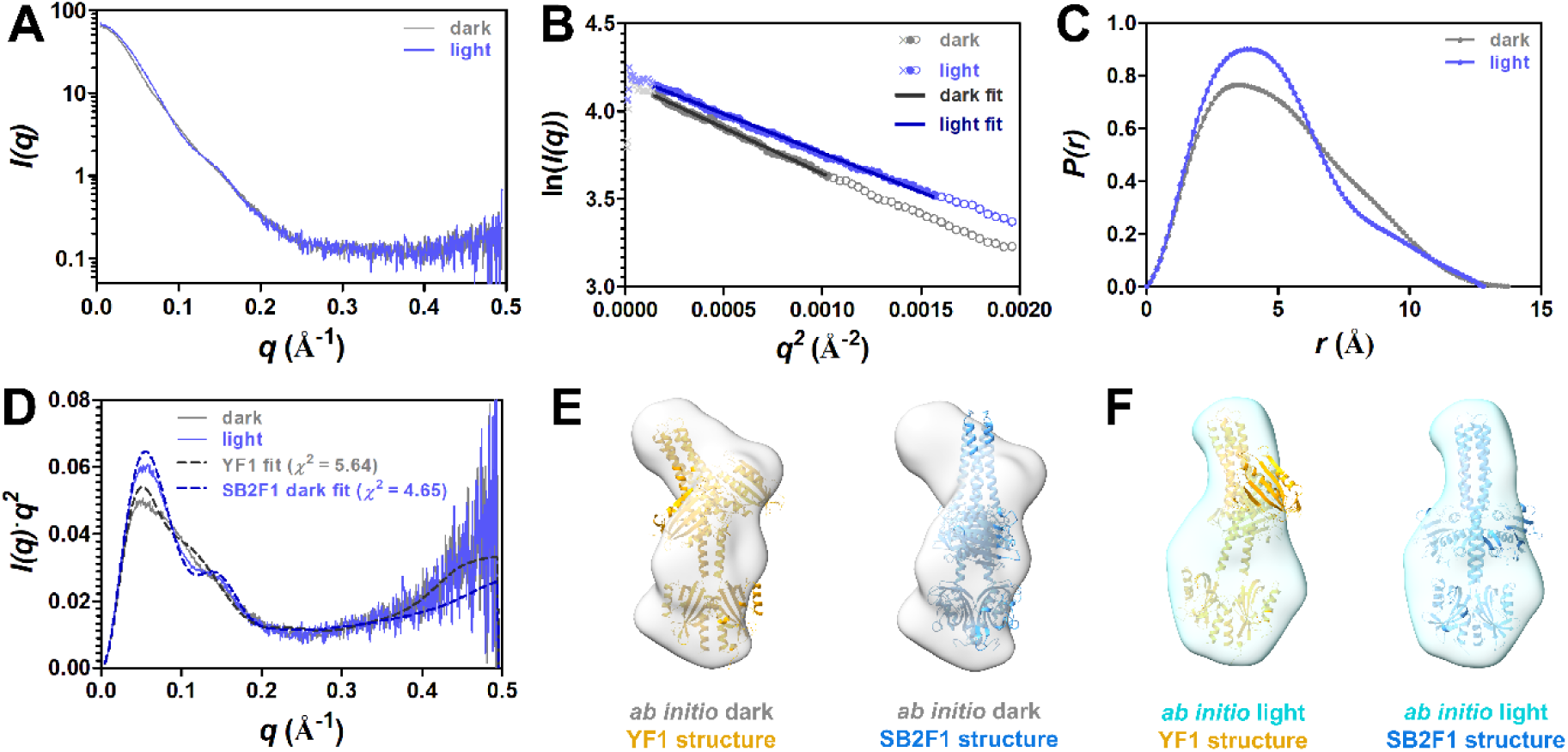
SAXS data obtained for ATP-bound SB2F1 in dark and light states. SAXS data of protein samples supplemented with 1.5 mM ATP, 2 mM MgCl_2_. (A) SAXS scattering curves, (B) Guinier plot for *q*×*R*g<1.3. Open symbols indicate data beyond the Guinier region. Data points in the low *q* region, represented by crosses, were excluded. Solid lines represent the Guinier fit. (C) Pair distribution function, *P(r)* (D) Kratky plot and Crysol-based model evaluation. The theoretical scattering curves (grey and blue dashed lines) calculated from the YF1 dark state structure (PDB-ID: 4GCZ, [29]) and SB2F1 dark state structure (PDB ID: 7A6P, [48] are shown. (E, F) Averaged and filtered *ab initio* models (transparent surface; grey: dark state, cyan: light state) superimposed with the YF1 dark state structure (PDB-ID: 4GCZ, [29]; orange cartoon) and SB2F1 dark state structure (blue cartoon). For clarity, FMN and ADP/ATP ligands are not shown. Density map shown at a contour level of 1σ, colored as indicated in the Figure. Further details are shown in Supporting Figure S7.

Overall, neither of the theoretical scattering profiles of the here determined crystal structures (irrespective of potential functional state or variant) fits the experimental scattering data recorded for ATP-bound SB2F1 in the dark state very well (*χ^2^* between 23.3-24.4; Supplementary Table S3). Interestingly, the theoretical scattering curve calculated for the YF1 dark-state structure provides a much better fit (*χ^2^* = 5.64; Figure 5, D, grey dashed line; Supplementary Table S3). The opposite holds for the light-state SAXS data. Here, the theoretical scattering profile calculated for all SB2F1 structures provide a reasonably good fit (*χ^2^* = 4.66 – 6.32; Supplementary Table S3) with the ATP-bound dark-state SB2F1 structure yielding the best fit (Figure 5, D, blue dashed line; Supplementary Table S3). In contrast, the YF1 structure exhibits a poorer fit (*χ^2^* =8.9; Supplementary Table S3).

The same can be seen in the *ab initio* envelope models calculated from the corresponding SAXS scattering data (Figure 5, E, F). Strikingly, the envelope calculated for the dark-adapted protein exhibits a marked kink whereas that determined for the light-adapted protein is largely symmetric. All SB2F1 structures much better fit the light-state envelope (exemplarily shown for the SB2F1 dark-state structure in Figure 5, F), while the kinked, highly asymmetric YF1 structure better covers the dark-state envelope (Figure 5, E). In conclusion, the SAXS data recorded for SB2F1 in dark- and light state reveal significantly larger structural changes than those seen in our crystal structures. All our structures (irrespective of assigned state) show a markedly better fit to the SAXS data recorded for the light state, which supports our earlier interpretation that they adopt a light-state like LOV-LOV dimer conformation.

## Discussion

### Structural analyses hint at intrinsic conformational equilibria

Irrespective of illumination, the arrangement of the LOV-LOV dimers in all our SB2F1 structures more closely resembles the light-state conformation of short-LOV protein structures than the dark-state conformation [43, 47, 55] (Figure 3, C and D, Supporting Table S2, Supporting Figure S5). Strikingly, exposure to light only induced overall modest structural changes within the SB2F1 crystal structures. By contrast, small-angle X-ray solution scattering revealed significantly larger structural changes induced by light (Figure 5). Whereas the solution scattering data acquired in the light agreed well with the SB2F1 structures, those in the dark did not but were instead consistent with the YF1 structure that shares with SB2F1 the identical DHp-CA module. This conundrum can be rationalized by assuming that LOV photoreceptors exist in a conformational equilibrium between light and dark states. Such intrinsic equilibria widely underlie function and allostery of many proteins [92, 93], not least in the model LOV2 domain of *Avena sativa* phototropin-1 (*As*LOV2), as revealed by nuclear magnetic resonance spectroscopy [94, 95]. Likewise, histidine kinases dynamically traverse their kinase-active and phosphatase-active states [22, 23]. We thus posit that such equilibria also govern the here designed SHKs and affect crystallization (Figure 6). Certain crystal lattice may thereby promote adoption of a dark-state like conformation, while other conditions select for a light-state like conformation. Consistent with this notion, the SB2F1-I66R dark- and light-state crystals (Table S1) all possess identical space groups (P3_2_21; trigonal), closely similar unit cell parameters and the same crystal packing (Supporting Figure S8) although they were grown in very different conditions. By contrast, YF1 crystallized in a hexagonal P6_5_22 space group, exhibited a pronounced kink in its structure (Figure 1)[29], with its LOV-LOV dimer displaying a short LOV dark-state-like dimer arrangement (Supporting Figure S9, Supporting Table S2).

**Figure 6:**
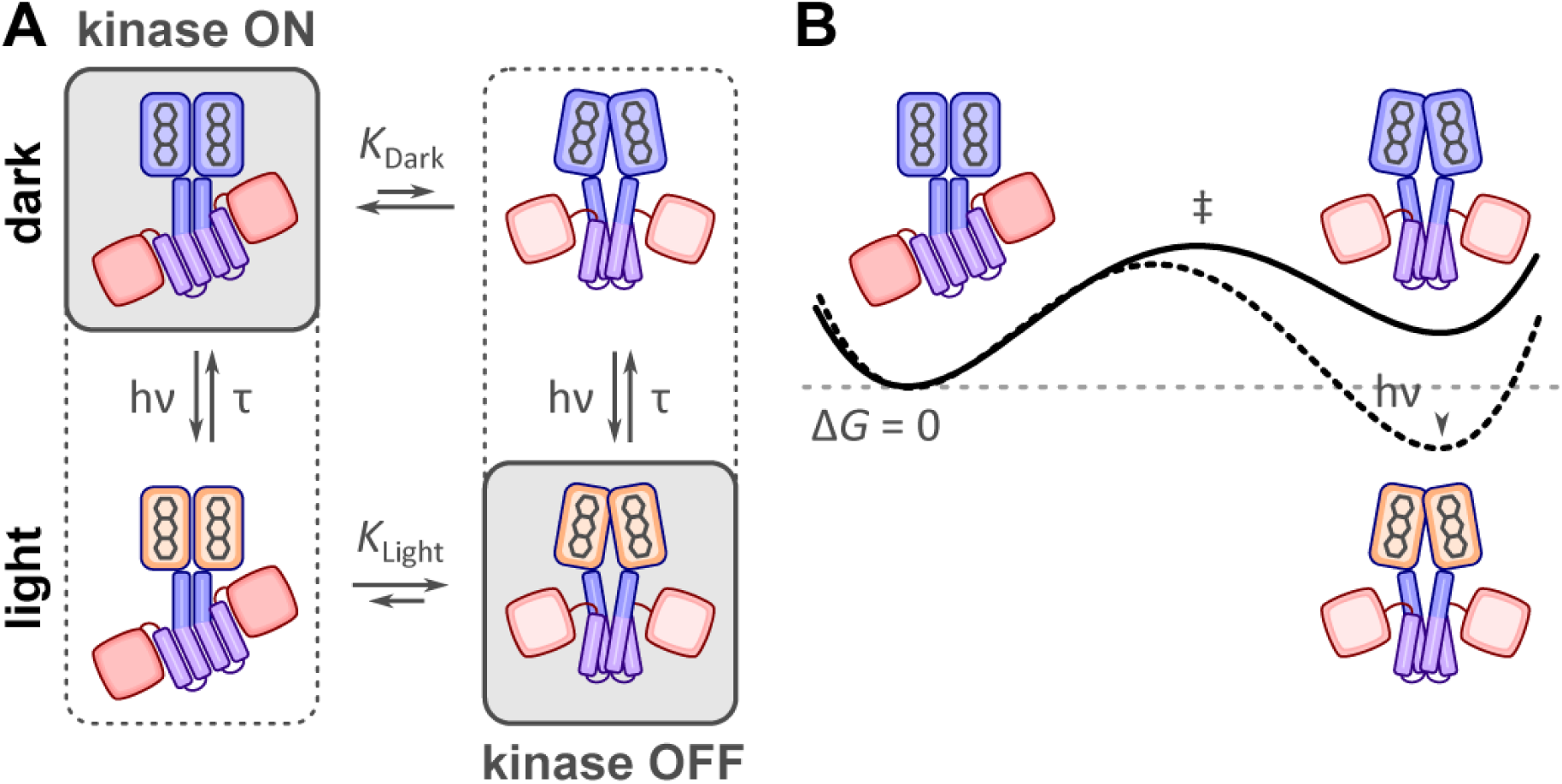
Conformational equilibria and signaling model of SB2F1. (A) Kinetic model of YF1/SB2F1 activation. In both the dark and the light-state, the YF1/SB2F1 SHK is in dynamic equilibrium between a kinase-ON and kinase-OFF state. Illumination (*hυ*) changes the equilibrium between the two states from *K*_dark_ in the dark to *K*_light_ and thereby modulates kinase activity. In terms of structure, this transition might be accompanied by the change from an asymmetric kinase-ON to a symmetric kinase-OFF dimer arrangement. The predominately populated states in darkness and under illumination are highlighted in grey. (B) Thermodynamic model of SB2F1 activation. Black and dashed lines denote the free-energy surfaces of the dark-state and the light-state, respectively. The kinase-ON state shown in the left has been arbitrarily assigned a Gibbs free energy of zero.

Importantly, functional assays revealed similar enzymatic activities and light responses in SB2F1 and YF1 (Figure 2). In darkness, they acted as net kinases and promoted RR phosphorylation, while upon exposure to blue light, they switched to net phosphatase activity. Based on these findings, SB2F1 and YF1, which share the same HK output module, may employ closely similar structural mechanisms for signal transduction. We therefore propose a light-induced transition from an asymmetric/kinked state (embodied by the YF1 dark-state structure) with net kinase activity (kinase ON) to a symmetric/straight state (exemplified by the SB2F1 structures) with net phosphatase activity (kinase OFF) (Figure 6). Support for this notion derives from SAXS (Figure 5), functional assays (Figure 2), conserved light-state like LOV-LOV dimer arrangements, a consistently large His154^…^ATP-γ-P distance (>19 Å) (not allowing for phosphor-transfer) in the straight SB2F1 structures and the observation of very similar (minor) differences between all of our SB2F1 structures (Supporting Figures S10 and S11).

### Light-dependent changes in Jα coiled-coil interactions drive asymmetry/symmetry transitions modulating kinase/phosphatase activity

We next analyzed global domain arrangements and local domain interfaces in the asymmetric/kinked YF1 vs. the symmetric/straight SB2F1 structures, as representatives of dark- and light states, respectively. To this end, we first evaluated the relative orientation of the two LOV protomers within the homodimeric HKs (Figure 7, A). For YF1, we observe a LOV-LOV crossing angle of around 84° compared to about 74° in SB2F1. Likewise, a significant difference between the A’α crossing angles of YF1 (32°) and SB2F1 (78°C) is observed, following the overall trend demonstrated for short LOV proteins [47, 55]. The rotational LOV-LOV dimer displacement goes along with a larger separation of the C-termini of the two LOV domains of about 2.1 Å in SB2F1 vs. YF1, highly reminiscent of the light-induced reorientation of the LOV-LOV dimer within the PpSB1-LOV protein and of light-induced structural transitions observed in solution for YF1 [96, 97]. Notably, the LOV domains rigidly connect to the adjacent Jα helices via their DIT/DVT motifs, a structurally conserved epitope at the C terminus of prokaryotic PAS domains [98]. Both in the SB2F1 and YF1 structures, the pertinent residues (D118-V119-T120 in SB2F1 and D125-I126-T127 in YF1) engage in several H-bonds between each other and a conserved tryptophan in the LOV domain (W103, YF1; W94, SB2F1) and thereby firmly anchor the Jα helices (Supporting Results 2; Supporting Figure S12). LOV domain rotation and slight pivoting apart of the DVT/DIT attachment points thus directly feed into the Jα helices and affect their relative orientation.

**Figure 7:**
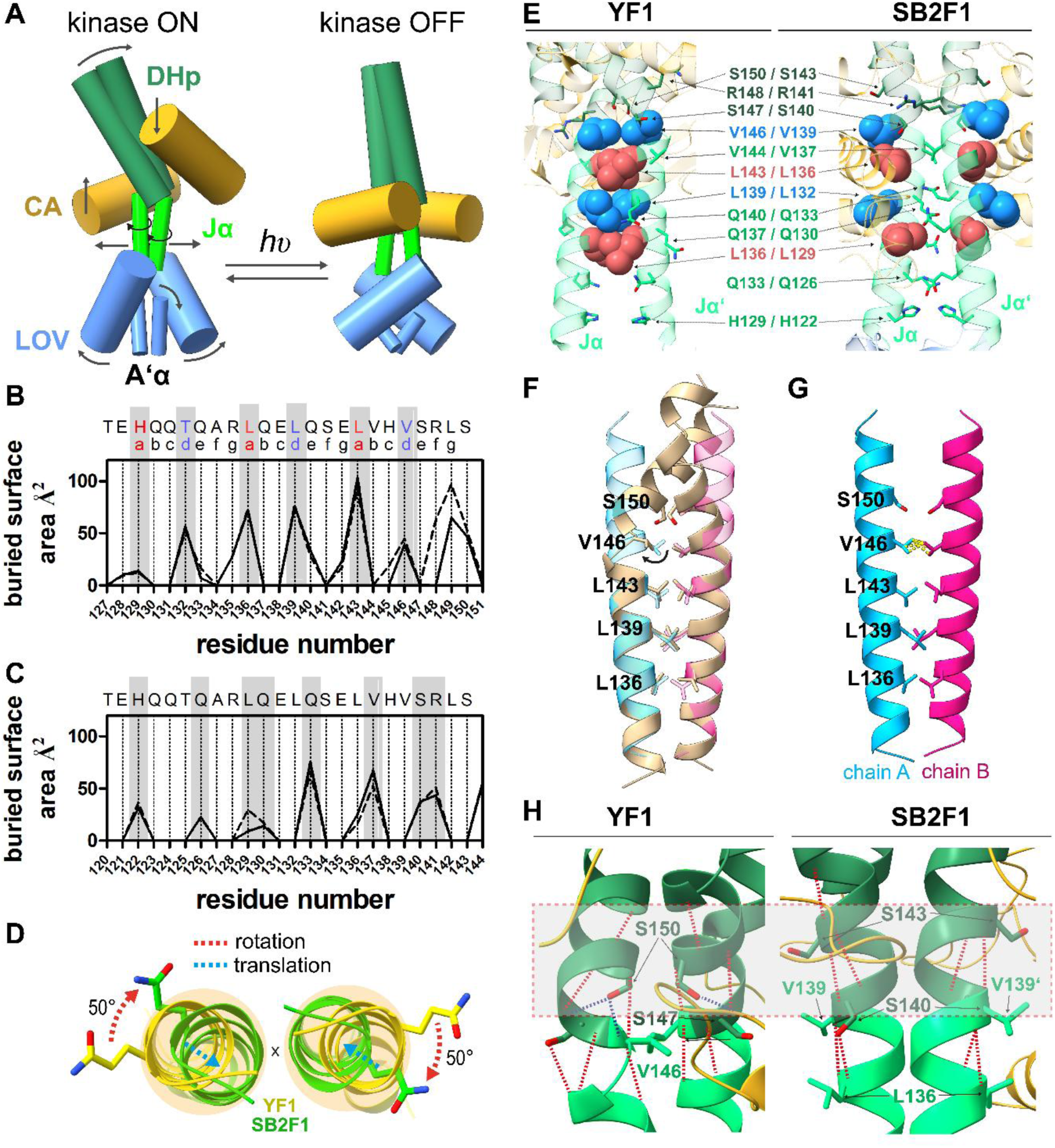
Interface analyses of SB2F1 and YF1. (A) Modular representation of asymmetric/kinked (YF1) and symmetric/straight (SB2F1) structures, representing SHK dark-(kinase ON) and light states (kinase OFF), respectively. Structural models generated by representing A’α-helices, LOV core domains, Jα-helices, DHP spine and CA domains by axes for which crossing angles were determined using ChimeraX. (B, C) Per-residue buried surface area derived from PISA [73] analysis for YF1 (B) [29] and the SB2F1 dark-state structure (C). Solid and dashed lines denote data obtained for the two chains of the dimer, respectively. Above panel B, the heptad register identified based on sequence and structure is given. (D) Rotational displacement of the Jα coiled coil viewed along the C2 axis (marked by a cross) from the LOV dimer towards the DHp spine. Helix rotation (red dashed line) and translation (cyan dashed line) inferred from LSQKAB superposition of the YF1 (yellow) and SB2F1 Jα helices (green). (E) Comparison of Jα–Jα′ coiled-coils with hydrophobic core residues as spheres, colored by heptad position (a: red, d: blue). Additional interface residues shown as sticks. (F, G) Structural model of a canonical (straight) YF1 Jα coiled coil, build using CCBuilder 2.0 [78], with individual chains (shown as cyan and pink transparent cartoon) superimposed on the kinked YF1 Jα coiled coil (F). Residues of the hydrophobic core are shown in stick representation. The black arrow marks the potential displacement of V146 induced by steric clashes in the canonical coiled coil structure. (G) Clashes between V146 of opposing chains in the canonical coiled coil model are highlighted (dashed yellow lines). (H) Jα–DHp junction in kinked (YF1) and straight (SB2F1) conformations. Key residues, including S150 (YF1) and S143 (SB2F1), highlighted in a grey box. H-bonds (<3.2 Å) shown as dashed lines (red: backbone; blue: side-chain). In all panels atom coloring as follows: C–domain/cartoon coloring, O–red, N–blue, P–orange.

Said Jα helices were proposed to form a short coiled-coil, arguably providing a rigid conduit along which the light signal propagates to the HK effector module [29]. Consistent with this model, two independent sequence-based coiled-coil prediction tools [75, 76] suggest the presence of a coiled-coil segment covering the Jα helix, in addition to two more within the DHp spine (Supporting Figure S13, A). The asymmetric YF1 structure indeed exhibits such a parallel Jα coiled coil (Supporting Figure S13, B) comprising two canonical heptad repeats (*abcdefg*) extending from H129-L149 (corresponding to H122-L142 in SB2F1), with the hydrophobic core formed by L136, L139, L143, and V146 within the *a* and *d* registers (Figure 7, B). The same coiled-coil element, with the same register, is also identified by the structure-based coiled-coil identification tool, Socket2 [77] and is also apparent in the repeating pattern of buried residues [73] inferred by PISA analysis (Figure 7, B). By contrast, in the symmetric SB2F1 structure the LOV protomers pull the Jα helices apart at their N-terminal ends and thereby preclude the formation of a canonical coiled coil. Notably, Socket2, even under the most relaxed packing cutoff, does not identify a coiled coil in the SB2F1 structure. As a corollary, the N-terminal portions of the Jα helices of SB2F1 run in parallel rather than being wound around another as in the YF1 structure. Intriguingly, the C-terminal halves of the Jα helices in the SB2F1 structure are sufficiently close to assemble into α-helical bundle, albeit with the heptad register shifted by one residue compared to the YF1 structure (identified by the repeating pattern of buried residues, Figure 7, C). For instance, residue V144 occupies the coiled-coil register *b* in the YF1 structure but the corresponding residue V137 in the SB2F1 structure falls within the register *a*. The altered register principally owes to a 50° rotation of each of the two Jα helices making up the coiled coil which amounts to a 100° relative reorientation of individual residue pairs within the juxtaposed helices (Figure 7, D).

As a consequence of this rotation, the hydrophobic core of the N-terminal segment of the Jα helix becomes solvent exposed in the SB2F1 structure (Figure 7, E) which contributes to a much reduced buried surface area and less stable Jα-Jα’ interface (solvation energy gain on complex formation SB2F1 Jα-DHp interface ∿ -19.3 kcal/mol compared to YF1: ∿ -24.9 kcal/mol [73]). Given the heptad periodicity of parallel dimeric coiled coils, consecutive residues within the constituent helices are angularly distributed in 100° increments around the helix spine. The differences in Jα helix orientation between the YF1 and SB2F1 and their impact on coiled-coil packing can thus be deemed roughly equivalent to insertion/deletion of single residues within the coiled-coil helices would have (also seen from the +1-residue shift of buried residues identified by interface analysis; Figure 7, C). In fact, ample biochemical data on YF1 and derivative variants indeed show that insertion of single residues within the Jα helix can lead to inversion of the response to light [99]. Collectively, these findings suggest that light promotes Jα-helix rotation, altering coiled-coil packing interactions, which likely translates into structural rearrangements in the DHP and CA domains, ultimately linked to catalysis (see below).

From the above comparison arises the question which structural features cause kinking of the YF1 Jα coiled coil, or conversely how unkinking is mediated and stabilized in the symmetric SB2F1 structure ─ both proposed here as intrinsic features of the YF1/SB2F1 signaling mechanism. To address this issue, we generated a structural model of a canonical (straight) YF1 Jα coiled coil and superimposed the individual chains onto the first two heptad repeats of the YF1 Jα structure (Figure 7, F, G). Clash analysis of the canonical model identified V146 as a primary site where steric clashes between opposing chains could destabilize the coiled coil (Figure 7, G), potentially initiating kinking. In fact, when we compare the kinked YF1 Jα coiled coil to the straight canonical one, we observe that in the kinked structure V416 is moved out of the interface (Figure 7, F; black arrow). As a consequence of kinking, the downstream S150 side chain is rotated to face into the hinge-region, forming H-bonding contacts to the backbone of S146 and V146 (Figure 7, H), thereby stabilizing the kink. In turn “unkinking” or straightening of the structure is a consequence of rational (torque-like) movement of the Jα coiled coil, which extends beyond the Jα element, with S150 (S143 in SB2F1) and V146 (V139 in SB2F) being rotated out of the coiled-coil interface (Figure 7, H). As a consequence, this also results in altered DHp-CA domain interactions which might contribute to stabilization of the symmetric/straight structure (Supporting Results 3; Supporting Figure S14 and S15).

As outlined in the results section, in the symmetric/straight dimer, the distance between the phospho-accepting His154 (corresponding to H161 in YF1) and the ATP γ-phosphate (Figure 3, A) is about 19-20 Å, which is too large to allow autophosphorylation, thus supporting the assignment of the symmetric/straight structure to the kinase OFF state. Note, however, that even the asymmetric/kinked YF1 structure does not fully capture the structure of the active kinase ON state, as the distance between the ATP molecule in the CA domain and the phospho-accepting His161 (corresponding to H154 in SB2F1) is with 12.8 Å still too large. Thus, additional conformational changes and/or a certain degree of flexibility are required to enable His161 autophosphorylation. We note that the CA domain connects to the DHp moiety via a linker of around 10 residues length that stands to grant exactly the type of motional freedom required to bring the γ-phosphate of the ATP into spatial proximity of the active-site histidine. However, further studies, that directly address intrinsic protein dynamics [100, 101], are needed to understand the importance of SHK flexibility and protein dynamics for catalysis.

In summary, we propose the following general model for photoactivation and signaling in SB2F1 and YF1 (Figure 7, A). In the dark, SB2F1/YF1 adopts an asymmetric/kinked dimer (similar to the YF1 dark-state dimer; as supported by our SAXS data). Such a conformation, combined with a general flexibility of the CA domains, allows autophosphorylation and phosphotransfer to FixJ (kinase ON state). Illumination triggers FMN-Cys adduct formation and results in a rotational movement of the LOV domains of the LOV-LOV dimer relative to each other. The resulting pivoting apart of the N-terminal ends of the Jα-helix is accompanied by a change of the overall Jα crossing angle and a torque-like rotational movement that translates into the subsequent DHp helical spine, resulting in a straight, more symmetric HK module conformation, with the phospho-accepting histidine moving away from the CA domain (kinase OFF state). Such a change is supported by our SAXS data and by the observation of light-state like LOV-LOV dimer arrangements seen for all of the here determined structures.

### Dimer asymmetry and flexibility drive signal transduction in LOV-SHKs and beyond

Apart from being relevant for designed SHKs like YF1 and SB2F1, the suggested signaling mechanism might also be realized by natural (LOV)-SHKs. In fact, DHp domain dimer asymmetry has long been associated with SHK signaling. One prevailing model suggests that SHKs exist as symmetric dimers with relatively straight α-helices in the DHp subdomain in the inactive/phosphatase state (kinase-OFF state) [102]. Upon activation (kinase-ON state), the dimer becomes asymmetric: one subunit adopts an autophosphorylating conformation, while the other engages in phosphotransfer with the RR. This transition involves bending of the DHp helices to accommodate the asymmetry. Hence, the kinase-ON state is associated with DHp bending/dimer asymmetry [102], while the kinase-OFF state is associated with a symmetric/straight dimer structure. A similar model has been suggested for the natural LOV-HK of *Brucella abortus*, which comprises an N-terminal LOV domain, a PAS domain, and a C-terminal HK module (DHp and CA domains) [11, 30]. Crystal structures revealed that the dark-state (kinase-OFF) LOV-PAS dimer (lacking the HK module) is straight and symmetric, while the light-state (kinase-ON) full-length LOV-PAS-HK dimer is highly asymmetric and kinked [30]. Solution scattering experiments and photoexcitation studies of the dark-grown LOV-PAS crystals further suggested a light-induced transition from a straight (kinase OFF) to a kinked asymmetric dimer (kinase ON) [30].

In addition, the suggested mode of signal-relay by rotational movement of a marginally stable coiled coil shares similarities with the signaling mechanism proposed for the periplasmic SHK BvgS of the whooping cough agent *Bordetella pertussis* [103], which together with its cognate RR BvgA regulates the expression of virulence factors necessary for infection. Each monomer of dimeric BvgS consists of two tandem periplasmic Venus fly trap domains, a transmembrane segment and a cytoplasmic PAS domain, linked via a helical connector that forms a parallel coiled coil in the dimer, to a HK output module. Intriguingly, in BvgS and related proteins, a DIT/DVT motif links the PAS domain to the coiled coil PAS-HK connector. Based on biochemical and reporter gene experiments it was shown that a flexible, rotationally dynamic, coiled coil favors the kinase ON state, while in the phosphatase-active (kinase OFF) state the coiled coil adopts a more rigid conformation that buries its hydrophobic interface [103]. This feature, although in reverse, is highly reminiscent of the model we suggest for YF1/SB2F1 signaling.

### Conclusions

Our study provides new insights into the signaling mechanism of engineered light-dependent LOV-HKs, highlighting that, similar to other LOV photoreceptors, they exist in a dynamic conformational equilibrium between light and dark states. This equilibrium, together with stabilizing interdomain and crystal contacts, might influence the structural outcomes observed in crystallization experiments, underscoring the importance of careful interpretation when assigning functional states based on full-length SHK structures. This is reminiscent of the discussion evolving for phytochrome based TCS, where structures of such proteins have yielded a number of partially conflicting conformational mechanisms for signal transduction (see [104], and discussion and references therein). The internal flexibility of the HK module (comprising the DHp and CA domains) reflects a key feature necessary for SHK function. This structural adaptability, complicates crystallization, but plays a crucial role in signaling and may lead to variations in observed conformations. Importantly, our data corroborate the central role of dimer asymmetry and HK module flexibility in SHK signaling, underscoring the importance of conformational diversity and dynamics in SHK function. As photoreceptor HKs serve as broadly applicable models for SHKs, our findings likely extend to many other members of this protein family, providing a framework for interpreting structural data and guiding the design of novel light-sensitive signaling systems.

## Abbreviations

CA: Catalytic/adenosine triphosphate (ATP)-binding domain
DHp: Dimerization/histidine phosphotransfer domain
FMN: flavin mononucleotide
HK: histidine kinase (HK)
LOV: light oxygen voltage
PAS: Per-ARNT-Sim
RR: response regulator

## Acknowledgements

This work was supported by grants from the German Federal Ministry of Education and Research (BMBF) (Project OptoSys, FKZ 031A16). We extend our sincere thanks to our former colleague, Dr. Joachim Granzin, for his invaluable expertise in crystal structure determination and his contributions to this work. AM and SSM acknowledge funding by the German Research Foundation (DFG; grant number: MO2192/4-2). The X-ray diffraction experiments were performed on the beamlines ID29 and ID30B, and SAXS data was collected at the beamline BM29 at the ESRF (Grenoble, France). We are grateful to local contact scientists at the ESRF for providing help in using the beamlines. Additionally, we are grateful to Dr. David von Stetten for the support on ID29S beamline for microspectrometry measurements.

## Author contributions

VA, AMS, UK and RBS conceived and designed the structural studies, functional assay, SAXS experiments, and spectroscopic measurements. UK designed and generated the SB1F1, SB2F1 variants. VA performed expression, purification, crystallization, production of mutants, and the dark recovery kinetic measurements. SM, AM and UK performed the functional assays. VA, and RBS collected X-ray data and performed structural analysis. VA, AMS, AM, UK, KEJ, and RBS analyzed and interpreted the data. RBS, and UK wrote the manuscript with contributions from all co-authors.

