## Supporting Information for "Dimer asymmetry in signaling of blue-light sensor histidine kinases"

Renu Batra-Safferling

Ulrich Krauss

Phone: +49 (0)921 / 55-7830

**The Supporting Figures and Tables are listed below in the order in which they appear in the main manuscript.**

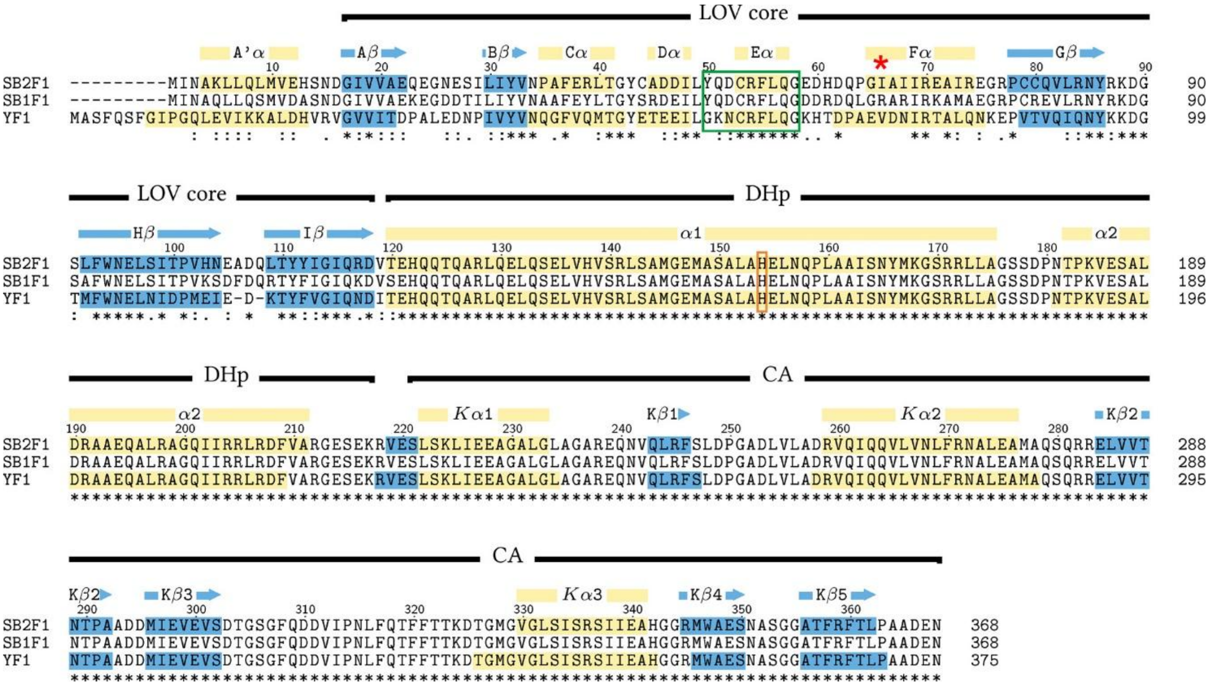

**Figure S1:** Multiple sequence alignment of SB1F1, SB2F1 and YF1. The green box marks the highly conserved GXNCRFLQG motif of LOV domains, which is altered to YXDCRFLQG in PpSB1-LOV, PpSB2-LOV and other *Pseudomonadaceae* short LOV proteins [1]. The red box marks His154, which is autophosphorylated. The red asterisk highlights the I66 position in SB2F1, which is mutated to Arg in SB2F1-I66R. SHK domain boundaries: LOV-domain, DHp and CA domain are labeled with bars above the alignment.

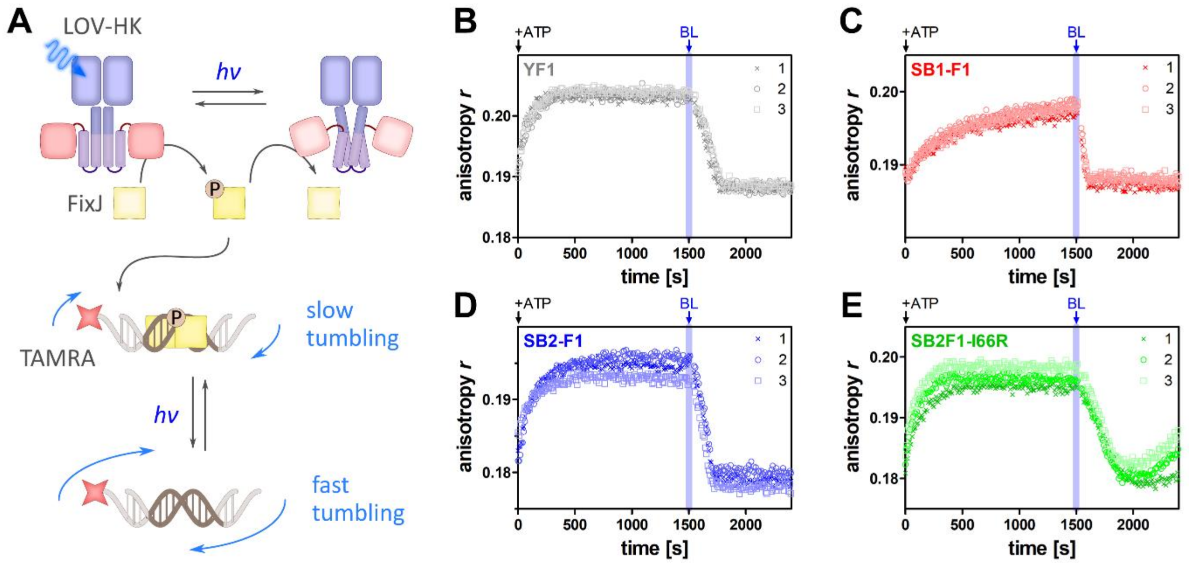

**Figure S2:** Histidine kinase and phosphatase activity of SB1F1, SB2F1, SB2F1-I66R and YF1 monitored by fluorescence anisotropy. A Scheme illustrating the assay principle [2]. All LOV-HKs possess net kinase activity in the dark phosphorylating the RR FixJ upon addition of ATP. Phospho-FixJ subsequently binds to its cognate operator sequence on a double-stranded DNA oligonucleotide labeled at its 5'-end with tetramethylrhodamine (TAMRA). Phospho-FixJ binding to the DNA target slows down the rotational tumbling of the labeled DNA fragment resulting in an increased TAMRA fluorescence anisotropy. Blue-light illumination (blue bar) enhances the phosphatase activity of the LOV-HK resulting in rapid dephosphorylation of Phospho-FixJ and unbinding from the labeled DNA fragment, hence rapidly decreasing the TAMRA fluorescence anisotropy signal. (B-E) Experimental results for SB1F1, SB2F1, SB2F1-I66R and YF1. All experiments were performed in triplicates (n = 3).

### Table S1: SB2F1 crystal structures: data collection and refinement statistics.

| Protein (State) | SB2F1 (Dark) | SB2F1 (Dark) | SB2F1 (illuminated) | SB2F1-I66R (Dark) | SB2F1-I66R (Light) |
|---|---|---|---|---|---|
| PDB ID | 8A3U | 8A6X | 8A52 | 8A7F | 8A7H |
| Beamline/Detector | ID29, ESRF (Grenoble, France)/DECTRIS PILATUS 6M-F (date 13-11-2015) | ID29, ESRF (Grenoble, France)/DECTRIS PILATUS 6M-F (date 13-11-2015) | ID30B, ESRF (Grenoble, France)/DECTRIS PILATUS3 6M (date 04-03-2016) | ID29, ESRF (Grenoble, France)/DECTRIS PILATUS 6M-F (date 21-06-2016) | ID29, ESRF (Grenoble, France)/DECTRIS PILATUS 6M-F (date 21-07-2016) |
| Wavelength (Å)/Monochromator | λ=0.97916/Silicon (1 1 1) | λ=0.97916/Silicon (1 1 1) | λ=0.97901/Silicon (1 1 1) | λ=0.97625 /Silicon (1 1 1) | λ=0.97625/Silicon (1 1 1) |
| Resolution range (Å), max. | 45.65 - 2.327 (2.510 - 2.327) | 45.43 - 2.452 (2.60 - 2.452) | 45.252 - 2.461 (2.639 - 2.461) | 45.345 - 2.71 (2.926 - 2.71) | 45.64 - 3.145 (3.336 - 3.145) |
| Space group | P $3_2$ 2 1 | P $3_1$ 2 1 | P $3_1$ 2 1 | P $3_1$ 2 1 | P $3_1$ 2 1 |
| Unit cell a=b, c (Å), α=β=90°, γ =120.0° | 138.72 49.35 | 138.79 96.08 | 138.25 94.99 | 138.53 96.89 | 139.42 89.88 |
| **Aimless/Staraniso** | | | | | |
| Total reflections | 277759 (12332) | 544226 (27231) | 504669 (12873) | 104836 (3851) | 244700 (12736) |
| Unique reflections | 13863 (693) | 27082 (1369) | 27319 (1387) | 16593 (831) | 12416 (621) |
| Multiplicity | 20.0 (17.8) | 20.1 (19.9) | 18.5 (9.3) | 6.3 (4.6) | 19.7 (20.5) |
| Completeness (%) spherical | 58.6 (14.6) | 68.7 (21.9) | 71.3 (19.4) | 56.1 (13.7) | 69.6 (21.6) |
| Completeness (%) ellipsoidal | 92.0 (63.3) | 95.5 (76.4) | 94.3 (59.4) | 92.1 (66.7) | 94.0 (68.2) |
| Worst diffraction limit after cut-off (Å) | 4,112 | 3,694 | 3.523 | 6,147 | 4,993 |
| Mean I/sigma(I) | 19.8 (1.5) | 20.2 (1.6) | 24.3 (1.4) | 13.4 (1.4) | 12.5 (1.5) |
| Wilson B-factor ($Å^2$) | 75.13 | 77.7 | 75.18 | 86.45 | 109.89 |
| CC (1/2) | 0.999 (0.480) | 0.999 (0.580) | 0.999 (0.493) | 1.0 (0.403) | 0.996 (0.483) |
| R-merge | 0.117 (2.260) | 0.122(2.366) | 0.102 (1.622) | 0.069 (1.078) | 0.163 (2.703) |
| R-meas | 0.120 (2.327) | 0.125 (2.429) | 0.105 (1.718) | 0.075 (1.216) | 0.168 (2.771) |
| R-pim | 0.027 (0.549) | 0.028 (0.543) | 0.024 (0.554) | 0.029 (0.555) | 0.038 (0.610) |
| Anomalous completeness (spherical) | 58.6 (14.8) | 68.7 (22.3) | 71.3 (19.8) | | |
| Anomalous completeness (ellipsoidal) | 91.9 (63.1) | 95.4 (6.5) | 94.3 (59.4) | | |
| Anomalous multiplicity | 10.4 (9.1) | 10.4 (10.1) | 9.6 (4.7) | | |
| CC(ano) | 0.922 (0.029) | 0.880 (0.026) | 0.945 (-0.004) | | |
| \|DANO\|/sd(DANO) | 2.220 (0.819) | 2.182 (0.787) | 2.187 (0.796) | | |
| **comment: phasing** | MSE-SAD phasing | MSE-SAD phasing | MSE-SAD phasing | MR with "8A6X" | MR with "8A6X" |
| **SAD-Phasing (SHELX-CDE)** | | | | | |
| fp/fdp | fp: -7.8705 fdp: 3.84174 | fp: -7.8705 fdp: 3.84174 | fp: -7.4025 fdp: 3.84046 | | |
| best shelxd solution | CC 52.83 CC(weak) 32.32 CFOM 85.15 | CC 52.41 CC(weak) 25.22 CFOM 77.64 | CC 55.81 CC(weak) 28.13 CFOM 83.94 | | |
| possible Se-sites (mol/asu) | 9 (1) | 18 (2) | 18 (2) | | |
| sites found (occ > 0.4) | 4 | 9 | 15 | | |
| contrast shelxe | none | 0.977 | 0.886 | | |
| rms coordinate differences to refined structure | not verified | 0.71 for 16 pairs | 0.66 for 16 pairs | | |
| after phase improvement and extension FOM | 0.481 | 0.574 | 0.589 | | |
| **Refinement (Phenix)** | | | | | |
| refined vs. | $F^+_{obs}/F^-_{obs}$ | $F^+_{obs}/F^-_{obs}$ | $F^+_{obs}/F^-_{obs}$ | $F_{obs}$ | $F_{obs}$ |
| Resolution range (Å) | 45.65 - 2.327 (2.423 - 2.327) | 44.61 - 2.452 (2.54 - 2.452) | 44.15 - 2.461 (2.549 - 2.461) | 41.07 - 2.71 (2.807 - 2.71) | 45.64 - 3.145 (3.257 - 3.145) |
| R-work | 0.2320 (0.4171) | 0.2259 (0.6222) | 0.2432 (0.5106) | 0.2283 (0.4655) | 0.2533 (0.4156) |
| R-free | 0.2797 (0.5910) | 0.2667 (0.6726) | 0.2952 (0.6977) | 0.249 | 0.3056 (0.4467) |
| coordinate error (max.-likelihood based) | 0.43 | 0.41 | 0.43 | 0.29 | 0.44 |
| Number of non-hydrogen atoms | 2891 | 5811 | 5784 | 5786 | 5756 |
| macromolecules | 2829 | 5687 | 5660 | 5662 | 5632 |
| ligands | 62 | 124 | 124 | 124 | 124 |
| Protein residues | 364 | 732 | 728 | 728 | 724 |
| RMS (bonds) | 0.004 | 0.008 | 0.005 | 0.005 | 0.005 |
| RMS (angles) | 0.64 | Jan-17 | 0.77 | 0.84 | 0.93 |
| Ramachandran favored (%) | 97.79 | 97.66 | 97.38 | 98.9 | 97.08 |
| Ramachandran outliers (%) | 0 | 0 | 0 | 0 | 0 |
| Clashscore | 5.78 | 14.29 | 8.84 | 14.70 | 14.99 |
| Average B-factor ($Å^2$) | 120.42 | 99.96 | 90.21 | 127.48 | 104 |
| macromolecules ($Å^2$) | 120.6 | 100.25 | 90.51 | 127.78 | 104.28 |
| ligands ($Å^2$) | 112.33 | 86.6 | 76.51 | 113.47 | 91.57 |
| Number of TLS groups | 5 | 11 | 10 | 9 | 11 |
| xtal conditions | 10% PEG8K, 0.1M Citrate 6.0, 0.2M NaCl, 0.1M Ammonium Bromide; 1mM ATP, 2mM MgCl2 | 10% PEG8K, 0.1M Citrate 6.0, 0.2M NaCl, 0.1M Ammonium Trifluoroacetate; 1mM ATP, | 6% PEG8K, 0.1M Na Citrate 6.15 0.2M NaCl; 1mM ATP, 2mM MgCl2 | 9% PEG8K, 0.1M Na-Citrate 5.7, 0.2M NaCl; 1mM ATP, 2mM MgCl | 18% PEG 3,3K, 0.1M Bicine 9.3, 0.2M LiSO4; 1mM ATP, 2mM MgCl |
| cryo conditions | + 15% PEG 8K, 20% Sucrose | + 15% PEG 8K, 20% Sucrose | 12% PEG8K, 0.1M Na Citrate 6.2, 0.2M NaCl, 30% PEG 200, 2mM MgCl2, 1.5mM ATP | + 20%PEG200, ATP/Mg | + 20%PEG200, ATP/Mg |
| Matthews coefficients $Å^3$/Da | 3.2 | 3.12 | 3.2 | 3.13 | 2.94 |
| solvent content % | 61.63 | 60.63 | 61.63 | 66.8 | 58.25 |

* *Statistics for the highest-resolution shell are shown in parentheses*

**Table S2: Root-mean-square deviation (RMSD) values between structural models.**

| **RMSD (Å) / *Nalign*** | SB2F1, dark PDB ID 8A6X | SB2F1-I66R, dark PDB ID 8A7F | SB2F1-I66R, light PDB ID 8A7H | YF1, dark PDB ID 4GCZ |
|---|---|---|---|---|
| SB2F1, dark PDB ID 8A3U | 1.0918 / 364[a] | | | |
| SB2F1, Illuminated PDB ID 8A52 | 0.6424 / 728[b] | | | |
| SB2F1-I66R, dark PDB ID 8A7F | 0.8972 / 728[b]<br>0.4561 / 236[c] | | | |
| SB2F1-I66R, light PDB ID 8A7H | 1.6327 / 724[b] | 1.9703 / 711[b] | | |
| YF1, dark PDB ID 4GCZ | 4.9704 / 302 | | | |
| PpSB2-LOV, dark PDB ID 7A6P | 3.2242 / 236[c] | 3.2268 / 234[c] | 3.086 / 235[c] | 1.7395 / 242[c] |
| PpSB1-LOV, light PDB ID 3SW1 | 1.0522 / 247[c] | 1.2189 / 251[c] | 0.8582 / 239[c] | 3.3312 / 242[c] |
| PpSB1-LOV, dark PDB ID 5J3W | 2.9959 / 241[c] | | | 1.5038 / 249[c] |

RMSD: Root mean-square deviation, a measure of the average distance between equivalent Cα atoms in two superimposed protein structures.
Nalign: Number of aligned residues included in the RMSD calculation after superposition.
a: residues from one protein chain; b: residues from dimer; c: residues from the LOV core domain of the dimer.
Worse fits with higher RMSD values are highlighted in orange boxes.
Calculations were performed using SSM superpose [3].

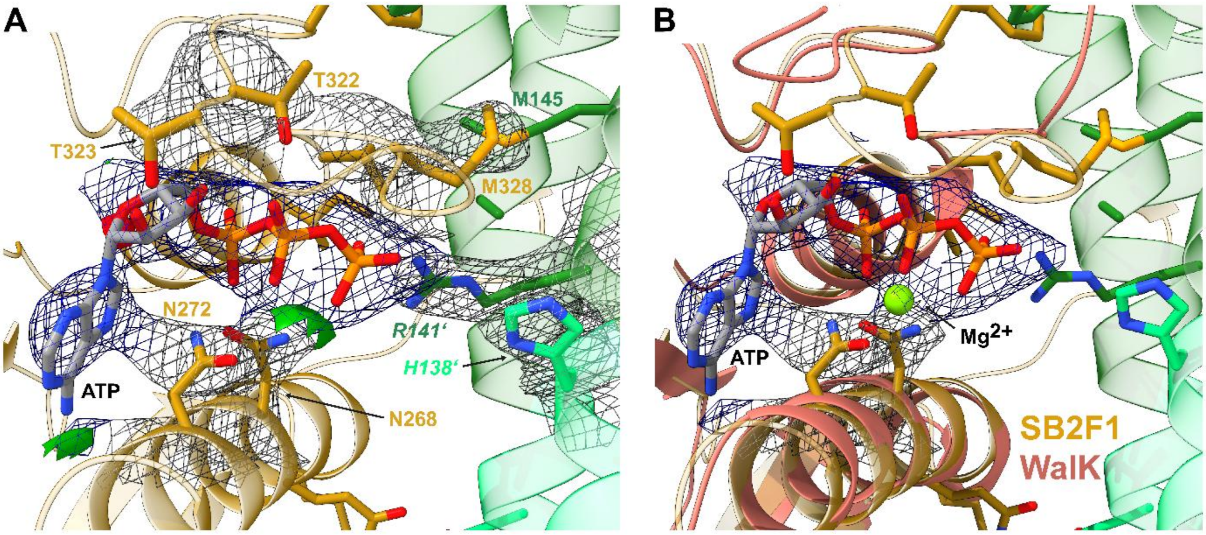

**Figure S3: SB2F1 ATP-binding CA domain and CA-DHp interface.** (A) SB2F1 dark-state structure and σ-A weighted 2mFo-DFc electron density map of the ATP molecule (blue mesh) and the CA-DHp domain interface residues (grey mesh) contoured at 1.0 σ. The CA domain of chain B and the DHp domain of chain B (dark/light helix in the back) and A (dark/light green helix in the front) are depicted. The mFo-DFc map is shown as colored green (3.0 sigma) and red (-3.0 sigma) surface 2 Å from the ATP molecule. (B) Superposition of the SB2F1 CA-DHp domain interface (CA domain, in gold, DHp domain in dark and light green) and the WalK CA domain structure (PDB ID: 3SL2, salmon; chain A). The $Mg^{2+}$ ion present in the WalK CA domain ATP binding site is shown as yellow green sphere. In addition, the 2mFo-DFc electron density map of the ATP molecule (blue mesh) and electron density around the N272, N268 side chains (grey mesh) contoured at 1.0 σ is shown.

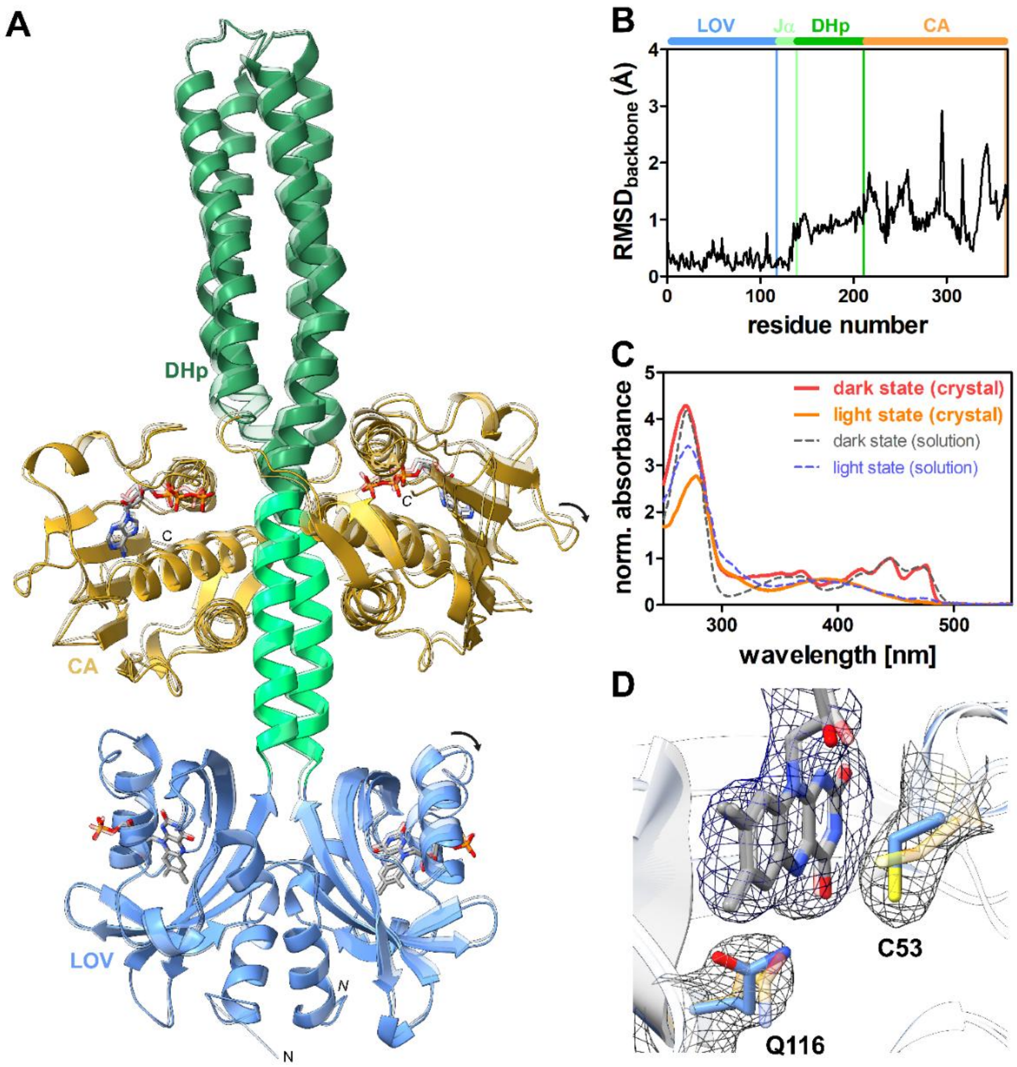

**Figure S4: Comparison of SB2F1 dark- and SB2F1 illuminated-state structure.** (A) Crystal structures of the SB2F1 illuminated-state structure (solid ribbon) superimposed with the SB2-F1 dark-state structure (transparent ribbon) with the domains colored as the follows: LOV – blue, DHp – light green and dark green (with the Jα-linker portion of the DHp α 1-helix in light green and the remaining DHp helical spine in dark green) and CA – gold. FMN and ATP are shown in stick representation with carbon in grey, oxygen in red, nitrogen in blue and phosphorous in orange. Structures superimposed via chain A of the LOV-LOV dimer (RMSD over backbone atoms, residues 1-363; 0.942 Å). (B) Residues-wise RMSD of the backbone atoms. Bars above the graph highlight domain boundaries (see Figure 3). (C) Single-crystal microspectrometry data of SB2F1 dark-grown crystals (red line) in comparison to SB2F1 solution dark- (dashed grey line) and light-state spectra (dashed blue line). Absorbance spectra recorded for the dark-grown SB2F1 crystal (red solid line), the illuminated dark grown crystal (solid orange line) and solution dark- and light state spectra (dashed grey and blue lines). All spectra were normalized to the 450 nm (dark state) and 390 nm (light state) absorbance band, respectively. Light-state (D) σ-A weighted 2mFo-DFc electron density map of the FMN chromophore, cysteine 53 and glutamine 116 of the SB2F1 illuminated-state structure contoured at 1.0 σ. The corresponding side chains and the FMN molecule are shown in stick representation with carbon atoms in blue, while the corresponding dark-state conformation is shown with carbon atoms in orange, respectively.

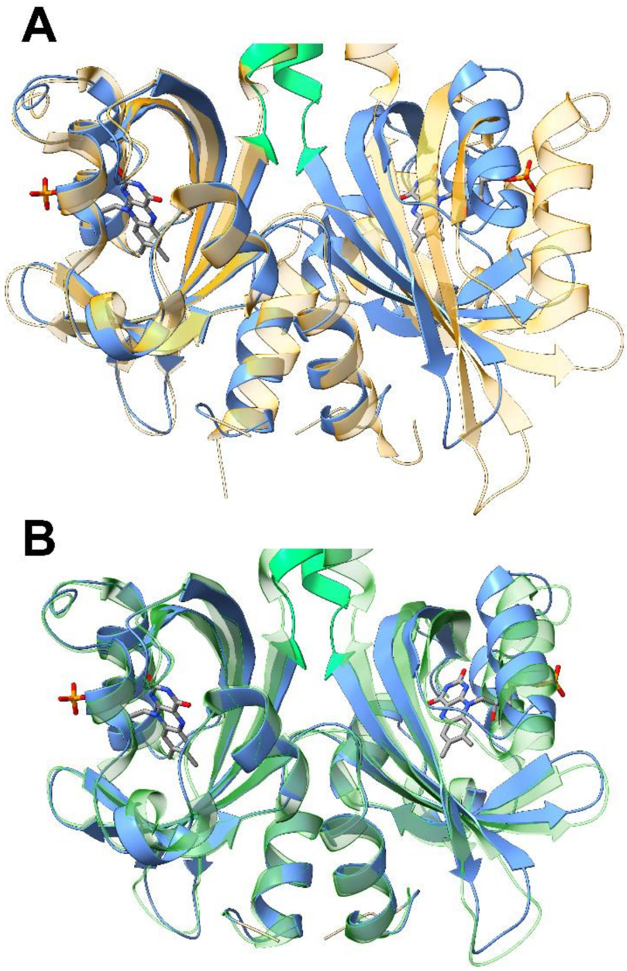

**Figure S5: SB2F1-I66R dark-state LOV-LOV dimer arrangement.** Superposition of the SB2F1-I66R dark state LOV-LOV dimer (blue solid ribbon) and (A) the PpSB2-LOV dark state structure (PDB-ID: 7A6P, [4])(transparent orange ribbon) (backbone RMSD over residues 1-118; 3.23 Å) (B) the PpSB1-LOV light-state structure (PDB-ID: 3SW1, [5])(transparent dark green ribbon) (backbone RMSD over residues 1-118; 1.05 Å). The FMN chromophore is shown in stick representation with carbon atoms in grey, oxygen in red, nitrogen in blue and phosphorous in orange.

**Supporting Results 1:**

As seen before in the SB2F1 illuminated-state structure, structural changes in the N-terminal sensory module include a minor rotation of the chain B in LOV dimer relative to chain A. This results in a decreased helix-crossing angle for the Jα helix portion of the DHp domain (12.8°) as compared to the dark state (15.6°). Such a change in angle could induce a torque like motion of the Jα helices, translating to a larger scale displacement of the remainder of the DHp domain. Consequently, the CA domains undergo a translational/rotational movement, resulting in a less parallel arrangement in the light-state dimer (crossing angle CA-CA; 14.6°) vs the dark state (8.8°). The most pronounced structural change appears to be the DHp domain displacement (Figure 4, A; Supporting Figure S6), which adopts a more bent-conformation in the light state. While the Jα-helix portion of the DHp domain can still be superposed quite well (average backbone RMSD over residues 119-139; 0.961 Å), the structures diverge beyond residue 137, reaching a maximal displacement for the loop connecting helix α1 and α2 and becomes minimal again at the DHp-CA junction (C-terminal end of the DHp helix α2 and the loop connecting the DHp to the CA domain; amino acids 212-218). This can be quantified by measuring the angle between the Jα-helix and the DHp domain of both chains of the dimer, yielding values between 4.9° (chain A) and 7.1° (chain B) for the dark state and 10.5° (chain A) and 11.9° (chain B) light-state structure.

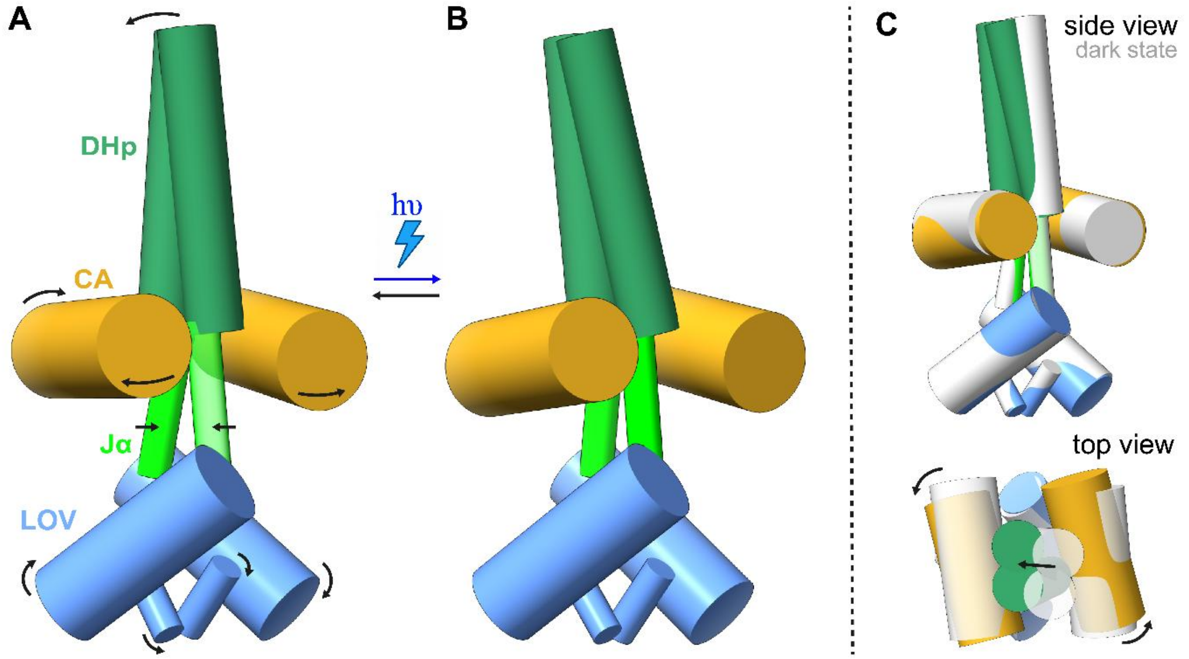

**Figure S6: Model depicting the global structural differences between SB2F1-I66R in the dark state (A) and light state (B).** The structures were modelled by fitting defined axis into the A'α-helix, the LOVcore, the Jα-helix, the DHp domain and the CA domain of both chains using ChimerX 1.9. Arrows in (A) mark movements of sub-structures (A'α and Jα-helix) and subdomains (LOV, DHp, CA) relative to the dark structure. (C) Side view and top view if superposed dark-state (grey) and light-state structures (colored as follows: as the follows: LOV – blue, DHp – light green and dark green (with the Jα-linker portion of the DHp α1-helix in light green and the remaining DHp helical spine in dark green) and CA – gold).

**Table S3: Summary of SB2F1 SAXS studies.**

| | **SB2F1 dark state / ATP** | **SB2F1 light state / ATP** |
|---|---|---|
| Calculated MW for monomer/dimer [kDa] | 42.8 / 85.6 | |
| SEC MW [kDa] | 117 | |
| DLS MW [kDa] | 99 | |
| SAXS MW from $I_0$ [kDa] | 64.3 | 66.7 |
| SAXS MW from Porod volume [kDa] | 80.0 | 78.8 |
| Crystal structure $R_g$ [Å] | 34 | |
| Guinier $R_g$ from SAXS [Å] | 39.6 ± 0.1 | 37.3 ± 0.1 |
| $D_{max}$ from SAXS [Å] | 137 | 128 |
| $\chi^2$ for the ATP-bound SB2F1 dark[2] | 23.332 | **4.655** |
| $\chi^2$ for the ATP-bound SB2F1 illuminated[2] | 24.347 | 5.428 |
| $\chi^2$ for the ATP-bound SB2F1-I66R dark[2] | 23.485 | 4.703 |
| $\chi^2$ for the ATP-bound SB2F1-I66R light[2] | 24.424 | 6.315 |
| $\chi^2$ for the YF1[1,2] | **5.637** | 8.902 |

1: limits $0.23 < q \times Rg < 1.25$ (dark state dataset), limits: $0.26 < q \times Rg < 1.29$ (light state dataset), [2:] $\chi^2$ value for the best fitting structure are highlighted in bold, respectively, [3:] PDB ID: 4GCZ [6]

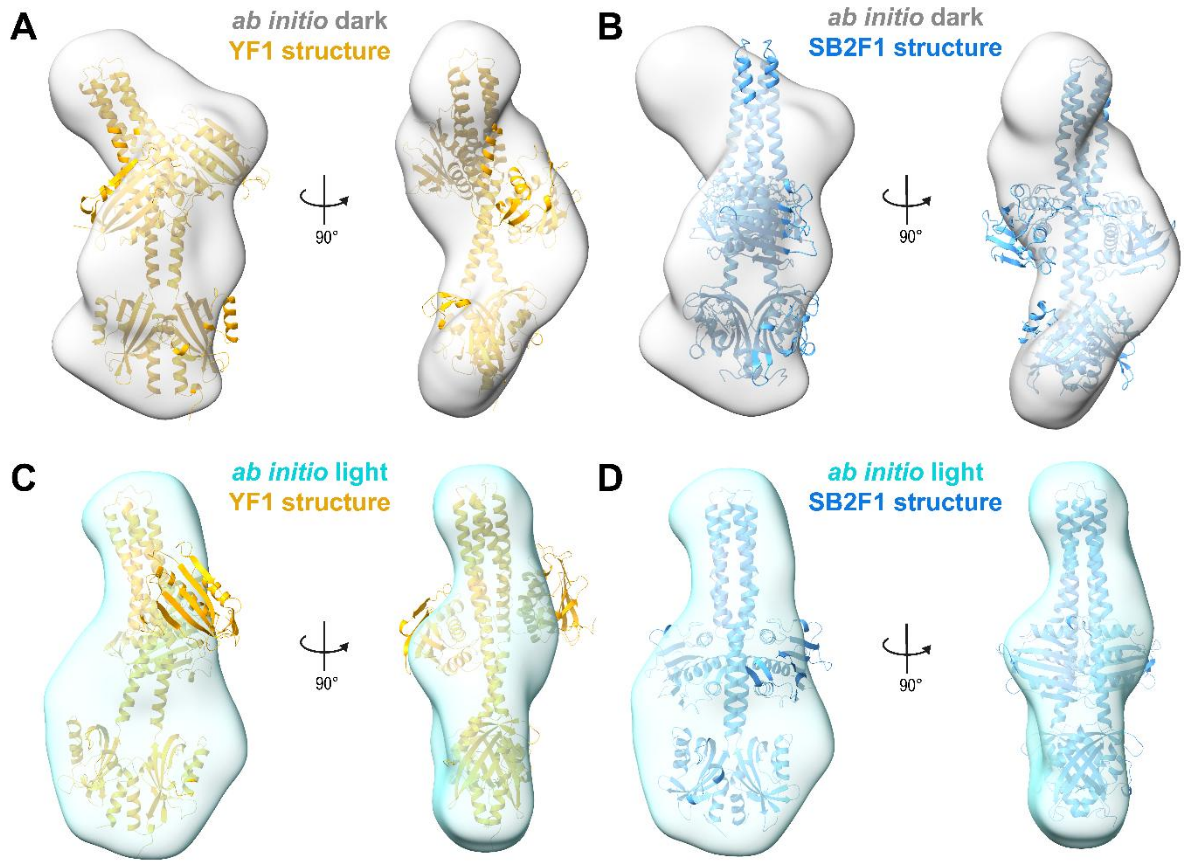

**Figure S7: *Ab initio* models evaluation**. Averaged and filtered *ab initio* models (transparent surface; grey: dark state, cyan: light state) superimposed with the YF1 dark state structure (PDB-ID: 4GCZ, [6]; orange cartoon) and SB2F1 dark state structure (blue cartoon). Maps contoured at 1σ. For clarity FMN and ADP/ATP ligands are not shown.

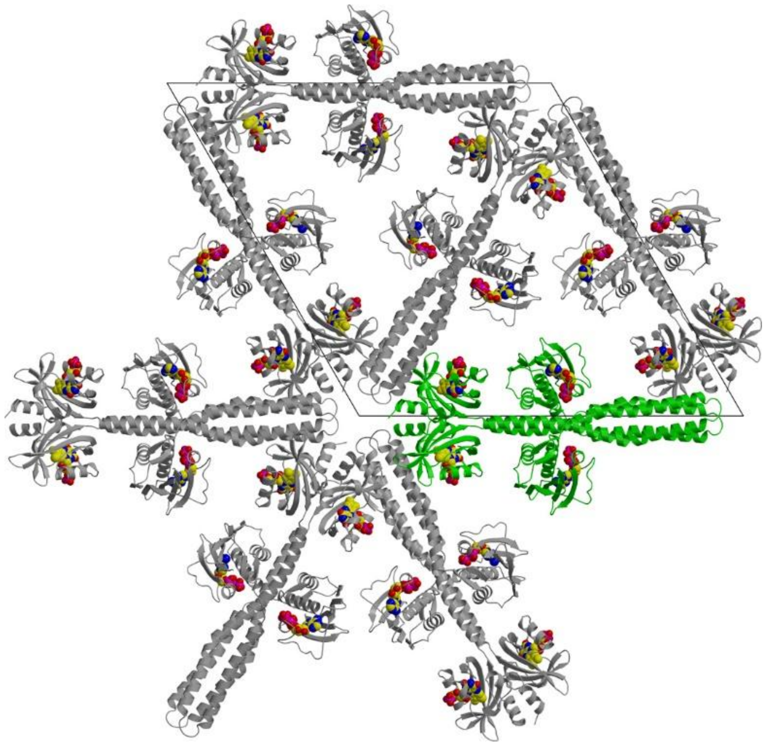

**Figure S8: Crystal packing in SB2F1 crystals.** Several symmetry-related SB2F1 dimers are depicted as ribbon representations in gray, with one dimer highlighted in green for clarity. Notably, the CA domains are positioned within solvent channels, allowing greater flexibility due to the absence of direct crystal contacts. Ligands ATP and FMN are shown as space-filling spheres, with carbon in yellow, oxygen in red, nitrogen in blue, and phosphorus in magenta.

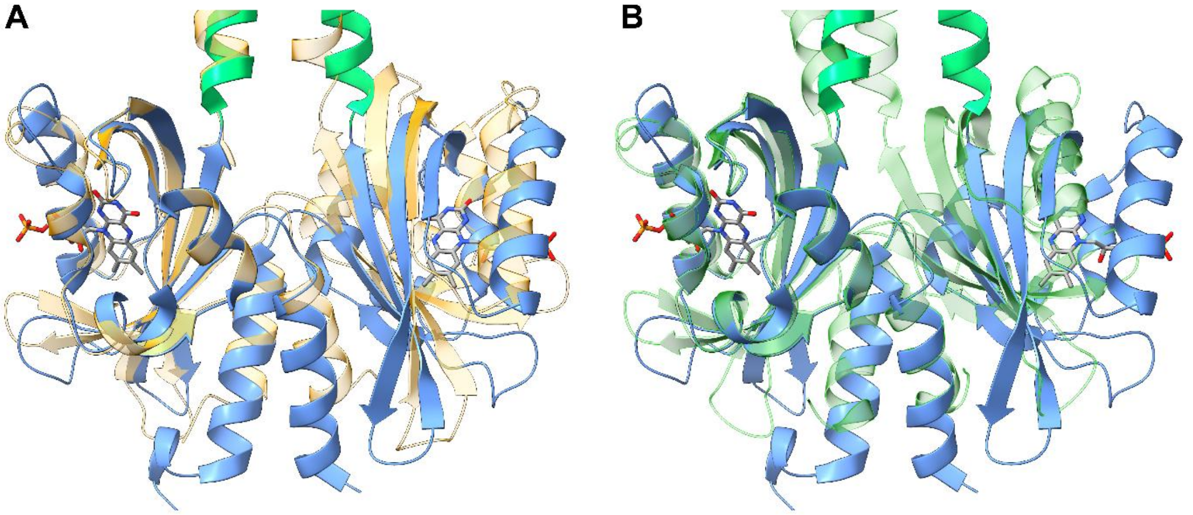

**Figure S9: YF1 dark-state LOV-LOV dimer arrangement.** Superposition of the YF1dark state LOV-LOV dimer (blue solid ribbon)(PDB-ID: 4GCZ; [6]) and (A) the PpSB1-LOV dark-state structure (PDB-ID: 5J3W, [7])(transparent orange ribbon) (3.37 Å backbone RMSD over residues 13-127 of YF1) (B) the PpSB1-LOV light-state structure (PDB-ID: 3SW1, [5])(transparent dark green ribbon) (6.96 Å backbone RMSD backbone RMSD over residues 13-127 of YF1). The FMN chromophore are shown in stick representation with carbon atoms in grey, oxygen in red, nitrogen in blue and phosphorous in orange.

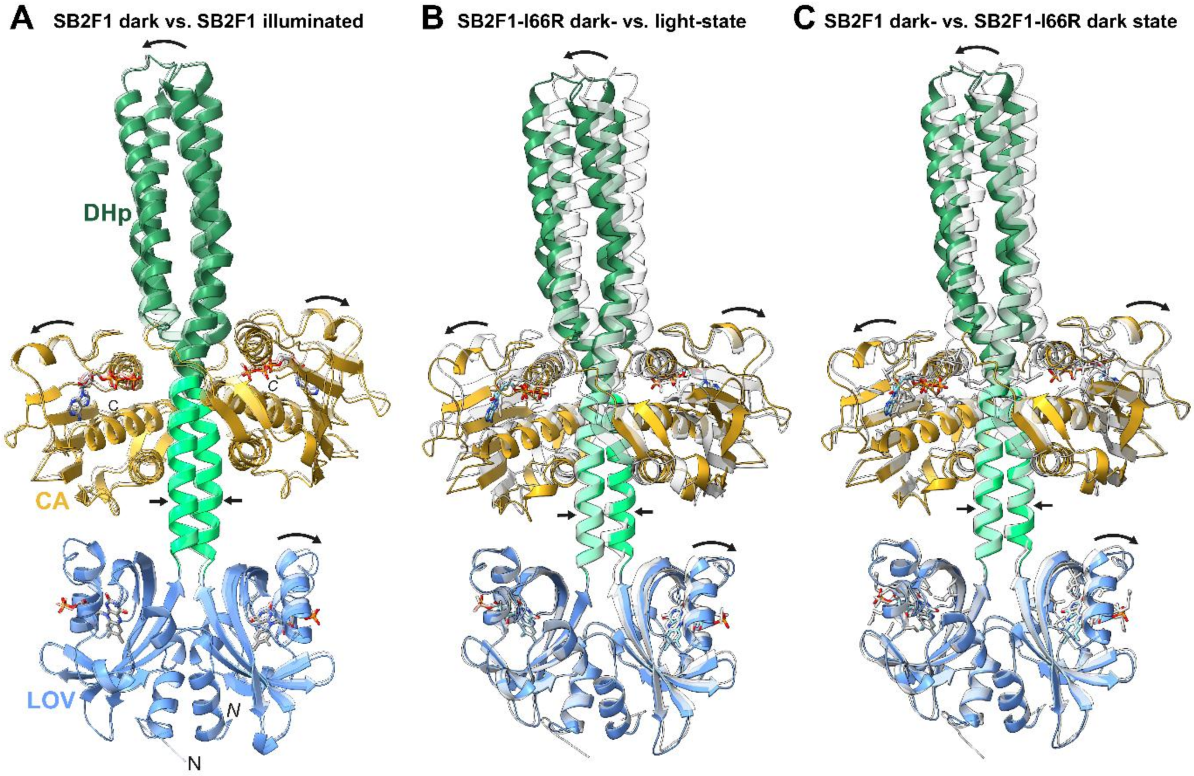

**Figure S10: Comparison of the light-dependent structural changes of SB2F1 and SB2F1-I66R and LOV-Jα-CA interface.** Superposition of (A) SB2F1 dark-state (grey transparent ribbon) and SB2F1 illuminated-state structure (solid ribbon, domain coloring), (B) SB2F1-I66R dark state (grey transparent ribbon) and SB2F1-I66R light-state structure (solid ribbon, domain coloring) and (C) SB2F1 dark-state (grey transparent ribbon) and SB2F1-I66R dark-state structure (solid ribbon, domain coloring). Domain coloring according to architecture as follows: LOV – blue, DHp – light green and dark green (with the Jα-linker portion of the DHp α1-helix in light green and the remaining DHp helical spine in dark green) and CA – gold. Arrows mark domain motions. The FMN chromophore and the ATP ligand are shown in stick representation with carbon atoms in grey, oxygen in red, nitrogen in blue and phosphorous in orange.

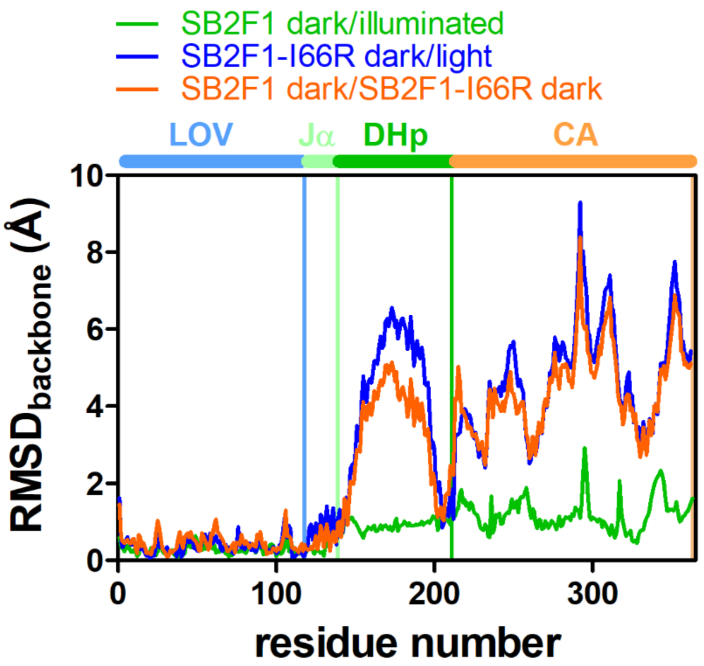

**Figure S11: RMSD-based comparison of the SB2F1 dark and SB2F1-I66R dark-/light-state structures.** Residues-wise RMSD of the backbone atoms for the comparison shown in Supporting Figure S10 A-C. Bars above the graph highlight domain boundaries (see Figure 3).

**Supporting Results 2:**

First, due to the different orientations of the two LOV domains in the SB2F1 and YF1 dimers, different inter- and intra-subunit interactions stabilize the respective LOV-LOV dimer structures (Supporting Figure S12, A, B), thereby likely contributing to the observed global asymmetry/symmetry. In the symmetric SB2F1 structure, inter-subunit interactions are present between the C-terminal end of the LOV-domain and the central β-scaffold of the opposite subunit (Supporting Figure S12, A), centered around K117, E96, S98 (K117-NH1···OE1-*E96'* salt-bridge, 3.74 Å; K117···*S98'* hydrogen-bond). The corresponding inter-subunit interactions are absent in the YF1 structure (Supporting Figure S12, B) with the corresponding residues N124, E105, N107 being solvent exposed (Supporting Figure S12, B). Instead, inter-subunit interactions are shifted towards the N-terminal interface present between the R24 (on the A'α-Aβ loop) and the β-scaffold of the opposite subunit (R24···*N107'*) (Supporting Figure S12, B).

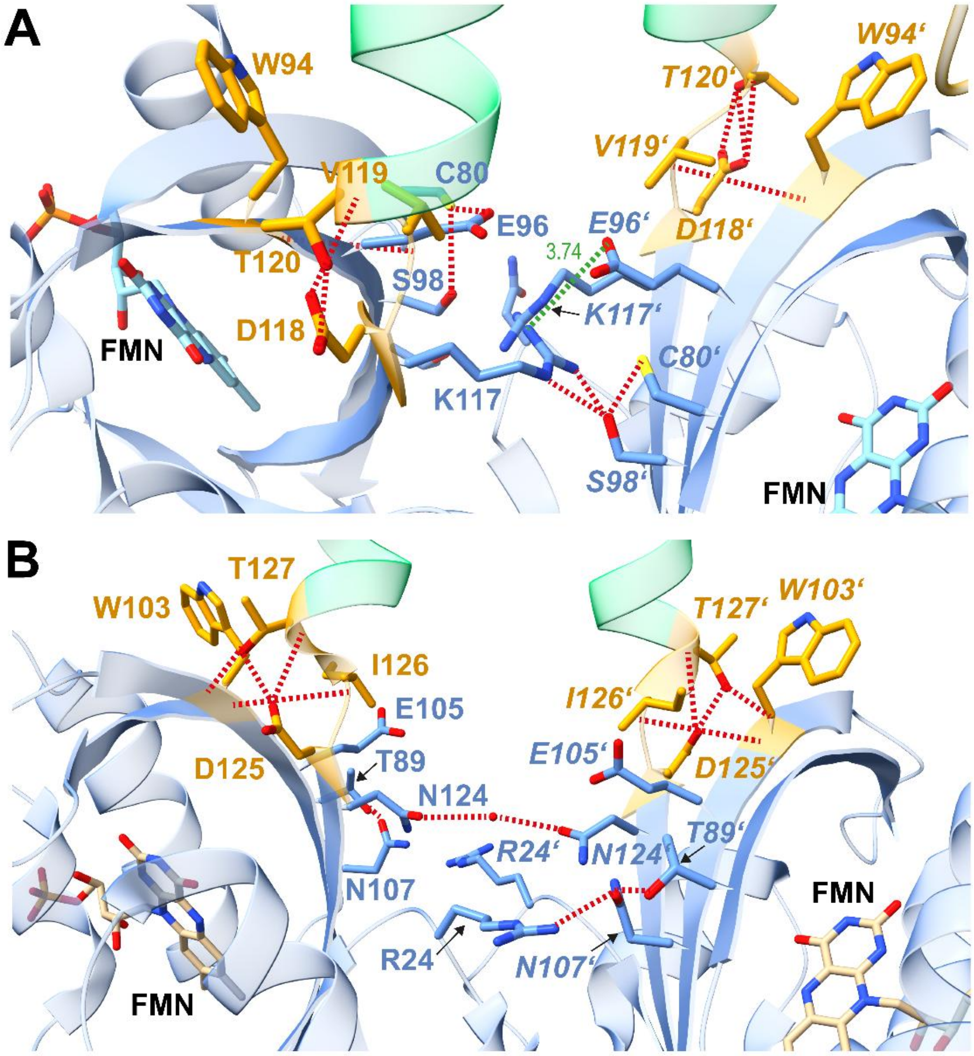

**Figure S12: Interface analyses of SB2F1 and YF1.** (A, B) LOV-LOV dimer interface of SB2F1 (A) and YF1 (B), shown as ribbon diagrams with LOV domains (blue), Jα helices (light green), and DVT (SB2F1) or DIT (YF1) motifs (orange). Key inter- and intra-subunit residues are shown as sticks with matching carbon colors. H-bonds (<3.2 Å) and salt bridges (<4.0 Å) are indicated as dashed red and green lines, respectively.

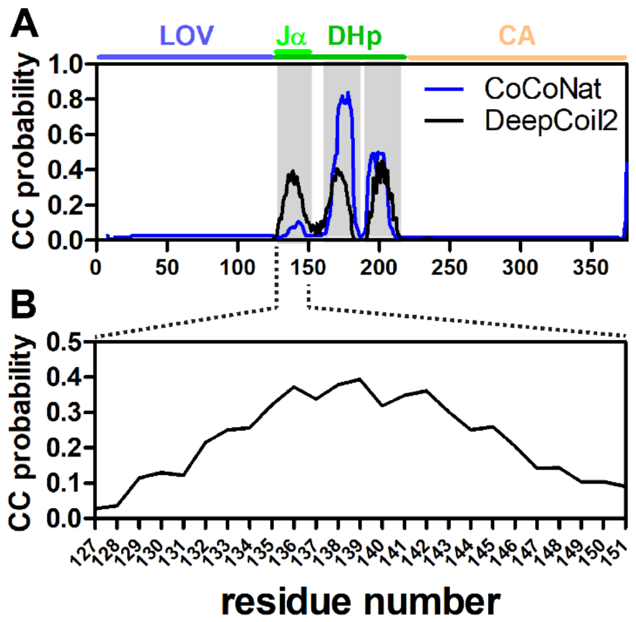

**Figure S13: Results of coiled-coil probability prediction for the YF1 sequence.** (A) The probability for the presence of canonical coiled-coil heptad repeat patterns was predicted based on sequence using the neural-network based coiled-coil prediction tools DeepCoil2 (data shown in black) [8] and CoCoNat (data shown in blue) [9]. Lines above the graph mark domain boundaries of the LOV, DHp (containing the Jα element) and CA domains. (B) Coiled-coil propensity, predicted by DeepCoil2 [8] for the Jα-helix segment of YF1.

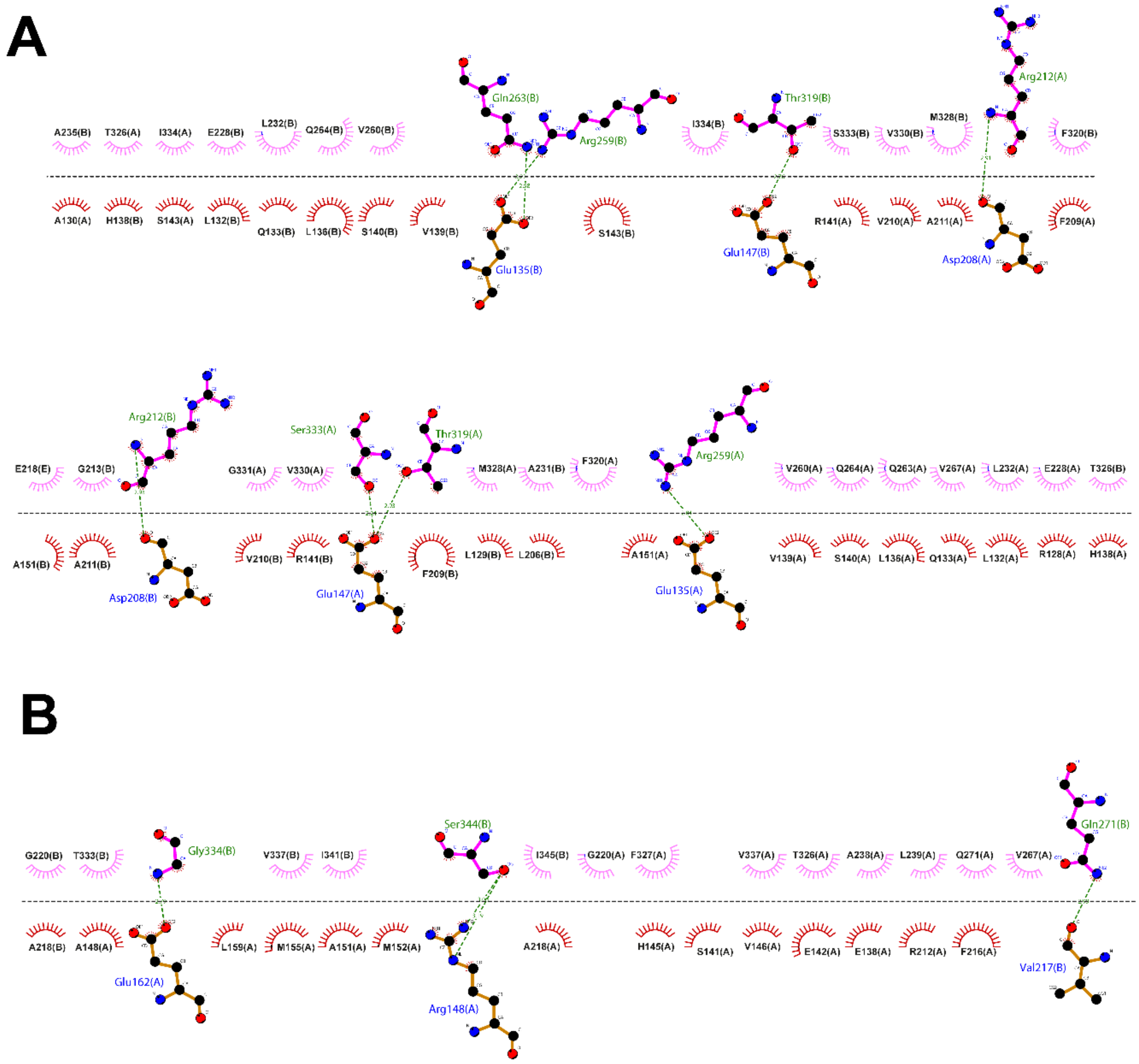

**Figure S14: DimPlot Analysis of the SB2F1 and YF1 Ja-DHp ··· CA domain interactions.** DimPlot analysis for (A) SB2F1 dark-state structure; DHp and CA domain defined as follows. DHp: 120-211 (phosphor-accepting His) CA: 212-363. And (B) YF1 (PDB: 4GCZ, [6]); DHp and CA domain defined as follows. DHp: 127-218 (phosphor-accepting His) CA: 220-370. All analyses were performed using the LigPlot+ v2.2. software tool [10].

**Supporting Results 3:**

“Unkinking” of the DHp domain also results in altered DHp-CA domain interactions which might contribute to stabilization of the symmetric/straight structure. Symmetric interactions between the Jα-DHp spine and the CA domains of both subunits include hydrophobic interactions between the exposed Jα core residues L132, L136, V139 and hydrophobic residues on the CA domain (mostly L232, V260). S143 (S150 in YF1), which in the kinked YF1 structure is buried within the kink (Figure 7, H), is rotated into the largely hydrophobic CA domain interface, pointing towards I334. In addition, H-bonding interactions are established between E135 (within the Jα helix) and Q263 and R259 within the CA domain. Further up the helical spine, H-bonding interactions are formed between Q147 (on the DHp domain), S333 and T319 within the CA domain (Supporting Figure S15).

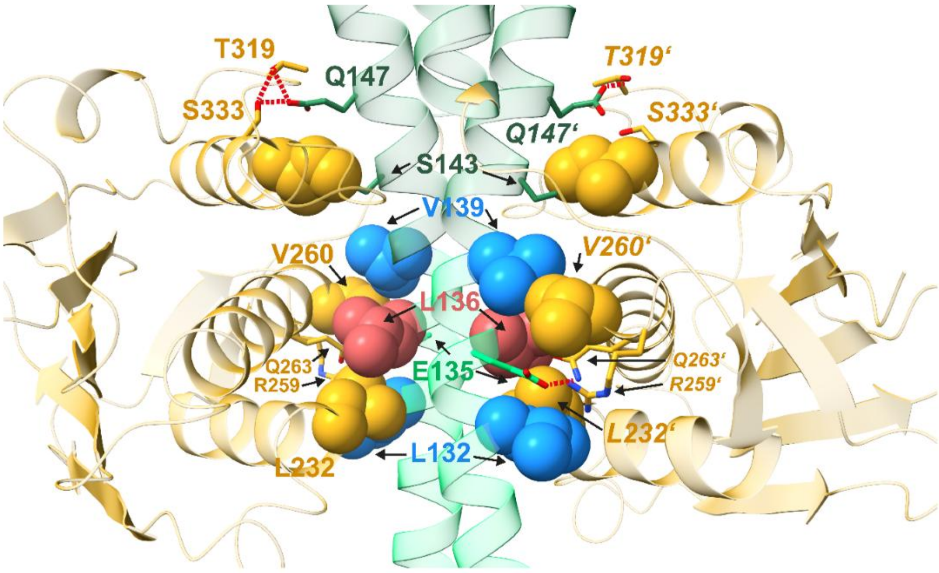

**Figure S15: SB2F1 DHp–CA interface.** Hydrophobic residues are shown as spheres and H-bonding/polar residues as sticks (H-bonds <3.2 Å; red dashed lines). In all panels atom coloring as follows: C–domain coloring, O–red, N–blue, P–orange.
